# Sonic Hedgehog Pathway Modulation Normalizes Expression of Olig2 in Rostrally Patterned NPCs with Trisomy 21

**DOI:** 10.1101/2021.10.19.465029

**Authors:** Jenny A. Klein, Zhen Li, Sanjeev Rampam, Jack Cardini, Amara Ayoub, Patricia Shaw, Angela L. Rachubinski, Joaquin M. Espinosa, Ella Zeldich, Tarik F. Haydar

## Abstract

The intellectual disability found in people with Down syndrome (DS) is associated with a decrease in white matter in the central nervous system. To study the mechanism of this myelination deficit, we differentiated two isogenic lines of induced pluripotent stem cells (iPSCs) derived from people with DS into brain-like and spinal cord-like neural progenitor cells (NPCs) and promoted a transition towards oligodendroglial fate by activating the Sonic hedgehog (SHH) pathway. In the spinal cord-like trisomic cells, we found no difference in expression of OLIG2 or NKX2.2, two transcription factors essential for commitment to the oligodendrocyte (OL) lineage. However, in the brain-like trisomic NPCs, OLIG2 is significantly upregulated and is associated with reduced expression of NKX2.2. We found that this gene dysregulation and block in NPC transition can be normalized by increasing the concentration of a SHH pathway agonist (SAG) during differentiation. These results underscore the importance of regional and cell type differences in gene expression in DS and demonstrate that modulation of SHH signaling in trisomic cells can rescue an early perturbed step in neural lineage specification in DS.

## Introduction

Down syndrome (DS) is a developmental disorder caused by triplication of human chromosome 21 (HSA21) (Lejeune et al., 1959). It is the most common genetic form of intellectual disability (ID) with a prevalence of 1 in 700 live births in the United States (Mai et al., 2019). The ID ranges from moderate to severe (Guéant et al., 2005), and while therapy and early intervention can improve outcomes in people with DS, the ID remains a significant factor impacting quality of life.

The exact etiology of the developmental brain defects in DS is unknown but is likely due to aberrant gene expression leading to several anatomical and physiological changes in the brains of people with DS. One prominent change is a well described white matter deficit that includes delays in the onset of myelination (Wisniewski and Schmidt-Sidor, 1989), and a lasting reduction in the density and organization of myelinated fibers (Ábrahám et al., 2012) (Olmos-Serrano et al., 2016). Gene expression analyses of mouse models and human brains indicate that a cell-autonomous change in OL maturation may represent a primary cause of the myelination deficit (Olmos-Serrano et al., 2016). As white matter is critical for propagating action potentials rapidly throughout the brain, disrupted myelination can affect the speed and timing of neural communication and thus contribute to the ID in DS.

Multiple genes on HSA21 are important for brain development including *DYRK1A, DSCAM, APP, OLIG1*, and critically for our study, *OLIG2*, which encodes a transcription factor essential for oligodendrocyte (OL) development (Zhou et al., 2001, 2000). Beyond specific HSA21 genes, trisomy 21 has widespread effects across the entire genome (Olmos-Serrano et al., 2016; Weick et al., 2013; Letourneau et al., 2014), including the alteration of sixty gene modules we identified using RNA isolated from the brains of people with DS (Olmos-Serrano et al., 2016). One of the modules significantly downregulated throughout development is comprised by genes that are important for myelination, specifically genes that are expressed in the maturing OL lineage (Olmos-Serrano et al., 2016). However, how these transcriptome changes translate to cellular consequences during OL precursor cell (OPC) specification, differentiation, or OL lineage maturation remains unknown.

OPCs are born at different times and in specific regions of the CNS in response to the morphogen Sonic hedgehog (SHH); OPCs in the spinal cord and ventral CNS are generated earlier than rostral brain and dorsal CNS OPCs (Cai et al., 2005; Fogarty et al., 2005; Kessaris et al., 2006; Pringle and Richardson, 1993). These regionally-specified groups of OPCs express different transcriptional profiles and differentiation trajectories (Bechler et al., 2015; Marques et al., 2018, Ghelman et al., 2021) indicating that intrinsic differences may result in functional specificity across regions. It is therefore important to consider spatial patterning in experimental design, especially in a condition like DS where gene expression is widely dysregulated both temporally and regionally (Olmos-Serrano et al., 2016). For instance, our prior work in the Ts65Dn mouse model of DS indicates that there are fewer OLs across development along with significantly fewer mature OLs in the corpus callosum (Olmos-Serrano et al., 2016) and in the dorsocortical spinal tract (DCST) (Aziz et al., 2019), a region of the spinal cord containing dorsally-derived OPCs (Tripathi et al., 2011). In contrast, the lateral funiculus (LF) region of the spinal cord, which is populated by ventrally-derived OLs (Tripathi et al., 2011), has more mature OLs throughout the lifespan and does not show a difference in the number of total OLs (Aziz et al., 2019). Thus, developmental origin is a critical feature to consider when investigating OL lineage cells in DS.

To investigate how these temporal and spatial aspects of early neural development are affected in DS, here we use two paired lines of isogenic induced pluripotent stem cells (iPSCs) derived from people with DS to investigate the effect of chromosome triplication and developmental origin on the early stages of the transition of NPCs towards an OPC fate. Each pair of isogenic lines is genetically identical except for the presence of a third copy of HSA21 in the trisomic lines ensuring that any changes observed can be attributed to the trisomy. Programming these cells’ development with different sequences of morphogens and small molecules, we measured changes in gene and protein expression associated with cellular differentiation. Using activation of the SHH pathway to prompt iPSCs to adopt a ventral and glial fate, we sought to determine how cellular differentiation is dependent on regional patterning, especially within the context of trisomy. We identified a SHH signaling based mechanism behind altered differentiation of the brain-like NPCs, suggesting that early region-specific perturbations in these cells may underlie later changes in OL differentiation and maturation in DS.

## Results

### Generation of PAX6+ NPCs with different regional characteristics

The two isogenic pairs used in our experiments consist of WC-24-02-DS-B (euploid, hereafter referred to as WC^eu^) and WC-24-02-DS-M (trisomic, hereafter referred to as WC^ts^) and ILD11#3 (euploid, hereafter referred to as ILD^eu^) and ILD1(2)-1 (trisomic, hereafter referred to as ILD^ts^). The cells were differentiated to NPCs and transitioned towards an OL fate following a published protocol with some modifications (Douvaras and Fossati, 2015) **(Figure 1A)**. PAX6, a classical marker for NPCs, was used to confirm the presence of NPCs at culture day 8 programmed either with Wnt-C59 (a Wnt pathway inhibitor, a driver of rostral CNS fate (Hermanto et al., 2019; Patapoutian and Reichardt, 2000)) or Retinoic Acid (RA, a driver of caudal CNS fate (Goldman and Kuypers, 2015; Lara-Ramírez et al., 2013)) **(Figure 1F,G,H)**. Confirming the efficacy of Wnt-C59, we found reduced mRNA expression of *Axin2*, a downstream target of the Wnt pathway (Jho et al., 2002), compared to cells cultured with RA (WC^eu^ 0.26x p=0.0027, WC^ts^ 0.46x p=0.0023; ILD^eu^ 0.45x p=0.066, ILD^ts^ 0.26x p=0.0015) **(Figure 1B).**

**Figure 1 –.**
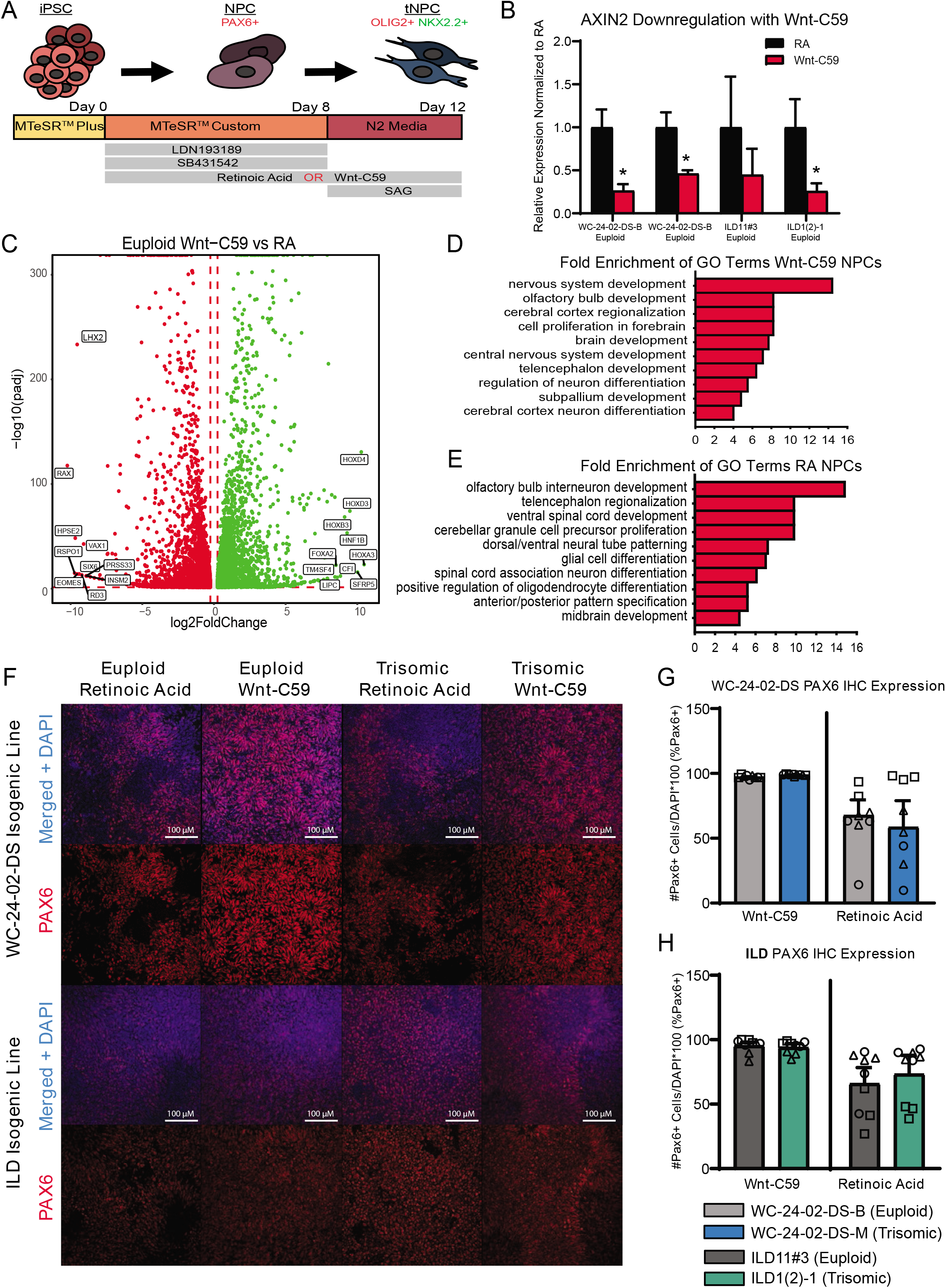
Generation of PAX6+ NPCs with different regional characteristics. **A)** Schematic of experimental design. **B)** Treatment of iPSC cultures with Wnt-C59 significantly decreases expression of *AXIN2*, a direct Wnt pathway target, in all cell lines used (WC^eu^, WC^ts^, ILD^eu^, and ILDts). **C)** 9,197 genes are differentially expressed in euploid NPCs differentiated with either RA or Wnt-C59. **D)** GO analysis identifies biological processes significantly enriched in Wnt-C59 NPCs relating to the rostral CNS. **E)** GO analysis identifies biological processes relating to the caudal CNS significantly enriched in NPCs treated with RA. **F)** Differentiation of the iPSCs with either RA or Wnt-C59 produce a pool of PAX6+ NPCs in both euploid and trisomic cultures. **G)** In the WC-24-02-DS isogenic pair, there are no significant differences either between genotype or condition in percentage of PAX6+ cells in culture, though the cultures treated with Wnt-C59 show less variability between differentiation experiments. **H)** The same pattern of PAX6 expression is repeated in the ILD isogenic pair. n=3,3 independent differentiation experiments with values from each individual well (3 per differentiation) overlaid. Data is shown as mean ± SEM. * p-value < 0.05 with specific values in text.

To assess whether our two patterning protocols resulted in NPCs with distinct spatial characteristics, we analyzed gene expression profiles of day 8 WC^eu^ NPCs exposed either to Wnt-C59 or RA using RNA-seq. In the euploid NPCs, we identified a total of 9,197 differentially expressed (DEX) genes between the two conditions with 4,622 being upregulated in the RA condition and 4,575 upregulated in the Wnt-C59 condition **(Figure 1C)**. Gene ontology (GO) analysis of the top 500 DEX genes in each condition resulted in a list of enriched biological processes including *cerebral cortex regionalization* (GO:0021796), *cell proliferation in forebrain* (GO:0021846), and *brain development* (GO:0007420) in Wnt-C59 treated cells, indicating that these NPCs showed regional characteristics of the rostral part of the CNS **(Figure 1D)**. In contrast, NPCs differentiated with RA included GO terms like *ventral spinal cord development* (GO:0021517), *spinal cord association neuron differentiation* (GO:0021527), and *midbrain development* (GO:0030901), supporting the conclusion that RA patterned the NPCs towards a more caudal fate **(Figure 1E)**. Similar analysis of trisomic NPCs also demonstrated similar regional characteristics **(Supplemental Figure 1)**. These results confirm that modulation of Wnt and RA signaling in isogenic human iPSC cultures can generate a regionally defined starting pool (“brain-like” vs. “spinal cord-like”, respectively) of euploid and trisomic NPCs for further differentiation studies.

Despite the changes in gene expression induced by the patterning molecules, both euploid and trisomic cells expressed PAX6 at similar levels with no significant difference between genotypes when differentiated with either Wnt-C59 (WC p=0.19, ILD p=0.99) or RA (WC p=0.99, ILD p=0.99) **(Figure 1G,H)**. However, we observed fewer PAX6+ cells in cultures patterned with RA compared to Wnt-C59. In the Wnt-C59 condition, 97.4% (± 0.3%) of WC^eu^ cells, 98.9% (± 0.2%) of WC^ts^ cells, 95.5% (± 2.7%) of ILD^eu^ cells, and 94.0% (±3.0%) of ILD^ts^ expressed PAX6. However, in the RA condition, only 67.8% (± 17.6%) of WC^eu^ cells, 58.6% (± 20.5%) of WC^ts^ cells, 66.1% (±12.4%) of ILD^eu^, and 73.5% (±14.5%) of ILD^ts^ expressed PAX6 **(Figure 1G,H)**. Despite these decreases, there are no significant differences between treatment condition (C59 and RA) within genotype (euploid: WC p=0.19, ILD p=0.20 and trisomic: WC p=0.19, ILD p=0.61). However, we also noted that the mean percentage of PAX6+ cells ranged widely in the RA condition compared to the Wnt-C59 condition. Using an F-test to quantify this variance, we found significant differences in variance between treatment conditions within genotype in the WC isogenic line (F(2,2)=1743.57, p=0.0011 for euploid and F(2,2)= 14193.23, p=1.4×10^−4^ for trisomic) and a trend towards differences in variance in the ILD isogenic line (F(2,2)=20.58, p=0.093 for euploid and F(2,2)=23.71, p=0.081 for trisomic). The reduced PAX6 expression as well as the higher variability between differentiation experiments in the RA condition indicate that treatment with Wnt-C59 results in a more homogeneous pool of NPCs.

### Trisomic spinal cord-like NPCs show no difference in transcription factor expression during transition towards OL fate

When NPCs patterned with RA were differentiated for four days with 1 μM of a Smoothened agonist (SAG) to activate the SHH pathway, they began to express the transcription factors OLIG2 and NKX2.2 **(Figure 2A)**, two SHH responsive genes that are essential for OPC differentiation (Zhou et al., 2001). We hereafter refer to these SAG-treated NPCs as transitional NPCs (tNPCs). We found no difference in the percentage of RA-patterned tNPCs positive for OLIG2 between genotypes. In the WC isogenic line 40.8% (± 9.2%) of WC^eu^ cells expressed OLIG2 compared to 42.5% (± 12.7%) of the WC^ts^ (p=0.92) **(Figure 2B)**. This expression pattern was observed again in the ILD isogenic line with 48.4% (± 4.4%) of ILD^eu^ cells expressing OLIG2 compared to 36.6% (± 10.7%) of the ILD^ts^ tNPCs (p=0.39). Similarly, no change was observed in the percentage of RA-patterned tNPCs positive for NKX2.2 **(Figure 2C).** 40.8% (± 17.7%) of WC^eu^ cells expressed NKX2.2 compared to 66.3% (± 9.6%) of WC^ts^ cells (p=0.29). Similarly, ILD^eu^ expressed NKX2.2 in 34.7% (± 3.2%) of the tNPCs while ILD^ts^ expressed NKX2.2 in 27.0% (± 14.4%) of the culture (p=0.65). Lastly, we found no significant difference between euploid and trisomic RA-patterned tNPCs co-expressing OLIG2 and NKX2.2 in either isogenic pair (p=0.42 WC-24-02-DS and p=0.90 ILD) **(Figure 2d; Supplemental Table 1**). Together, these data indicate that in spinal cord-like tNPCs, trisomy 21 does not affect the percentage of cells positive for OLIG2 or NKX2.2 following SHH activation.

**Figure 2 –.**
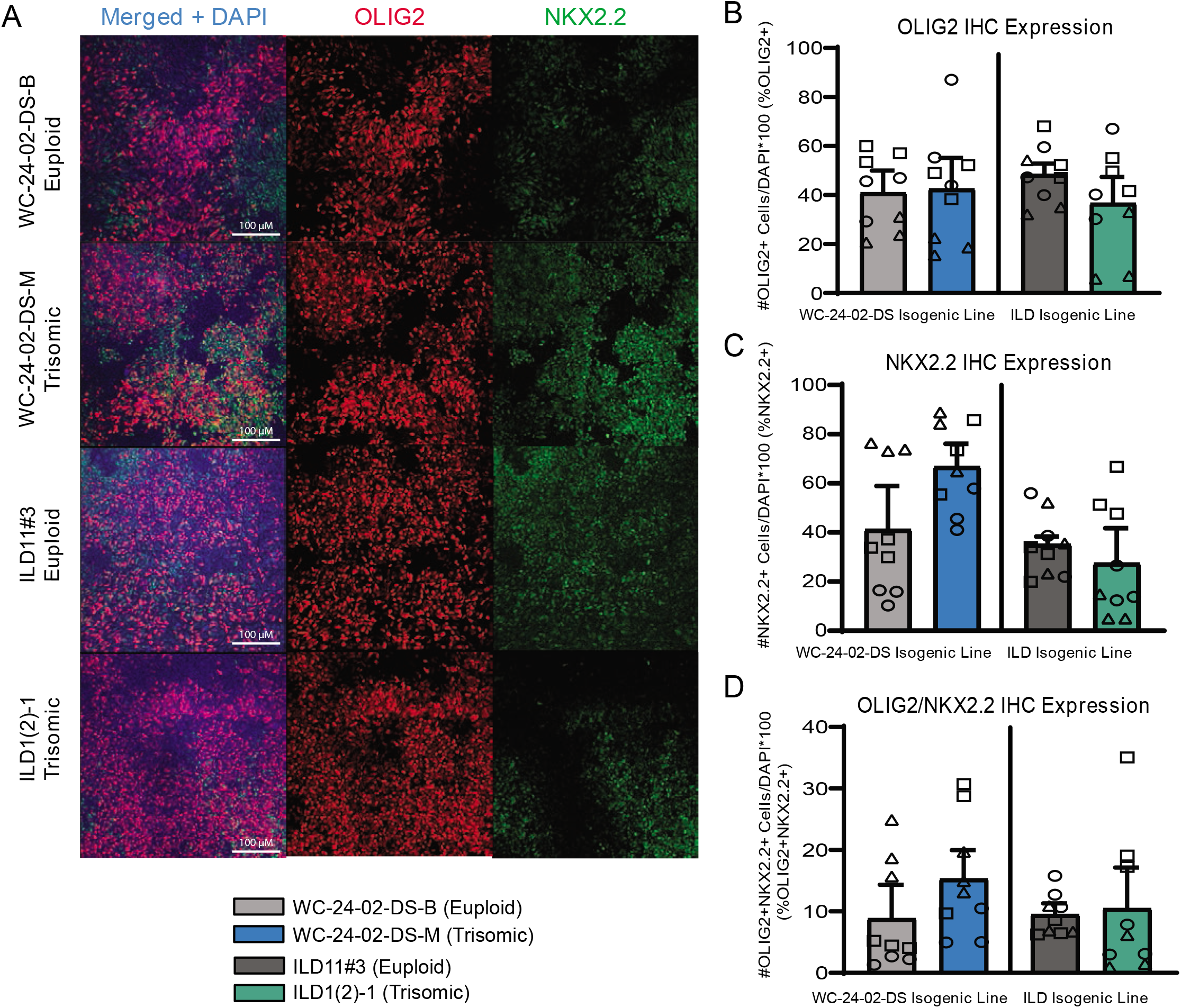
Trisomic spinal-cord like NPCs show no difference in transcription factor expression during transition towards OL fate. **A)** Both isogenic pairs of cells lines (WC-24-02-DS and ILD) differentiated with RA express OLIG2 and NKX2.2 in tNPCs after four days of treatment with SAG, a SHH pathway agonist. **B)** In both isogenic pairs, there are no significant differences in OLIG2 expression in the trisomic line (WC^ts^ and ILD^ts^) compared to their euploid controls. **C)** In both isogenic pairs, there is no significant difference in expression of NKX2.2 in the trisomic lines (WC^ts^ and ILD^ts^) compared to euploid. **D)** There is no difference in percentage of cells co-expressing OLIG2+ and NKX2.2+ between euploid and trisomic lines in either isogenic pair, n=3,3 independent differentiation experiments for all IHC measurements with values for individual wells (3 per differentiation) overlaid. Data is shown as mean ± SEM.

### Rostral patterned tNPCs show an altered response to SHH activation

When NPCs patterned with Wnt-C59 were treated with 1 μM SAG for four days, they also started to express OLIG2 and NKX2.2 **(Figure 3A)**. However, in contrast to RA patterning, there were significant differences in tNPC gene expression between genotypes. Specifically, the trisomic tNPCs expressed significantly higher levels of OLIG2 in both isogenic lines **(Figure 3B).** In the WC line, 11.4% (± 2.9%) of the WC^eu^ tNPCs expressed OLIG2 compared to 29.2% (± 3.4%) of the WC^ts^ tNPCs (p=0.018). In the ILD line, only 7.8% (± 0.7%) of the ILD^eu^ tNPCs expressed OLIG2 while 51.0% (± 2.8%) of ILD^ts^ tNPCs expressed OLIG2 (p=0.0028). In contrast, the number of tNPCs expressing NKX2.2 protein was decreased **(Figure 3C)**. The results in the WC line indicate that 15.4% (± 5.6%) of the euploid (WC^eu^) tNPCs expressed NKX2.2 while only 3.7% (± 0.4%) of the WC^ts^ tNPCs expressed NKX2.2. However, these differences were not statistically different (p=0.17) due to a high degree of variability. Similarly, 10.0% (± 3.7%) of the ILD^eu^ tNPCs expressed NKX2.2 while only 5.3% (± 1.9%) of ILD^ts^ tNPCs expressed NKX2.2 (p=0.34). We detected very few tNPCs co-expressing OLIG2 and NKX2.2 in either line or genotype, which was not surprising based on the early stage of differentiation. Nevertheless, we found no significant difference in OLIG2+/NKX2.2+ tNPCs between euploid and trisomic lines in either isogenic pair in the Wnt-C59 condition (p=0.30 WC-24-02-DS and p=0.57 ILD) **(Figure 3D; Supplemental Table 1)**.

**Figure 3 –.**
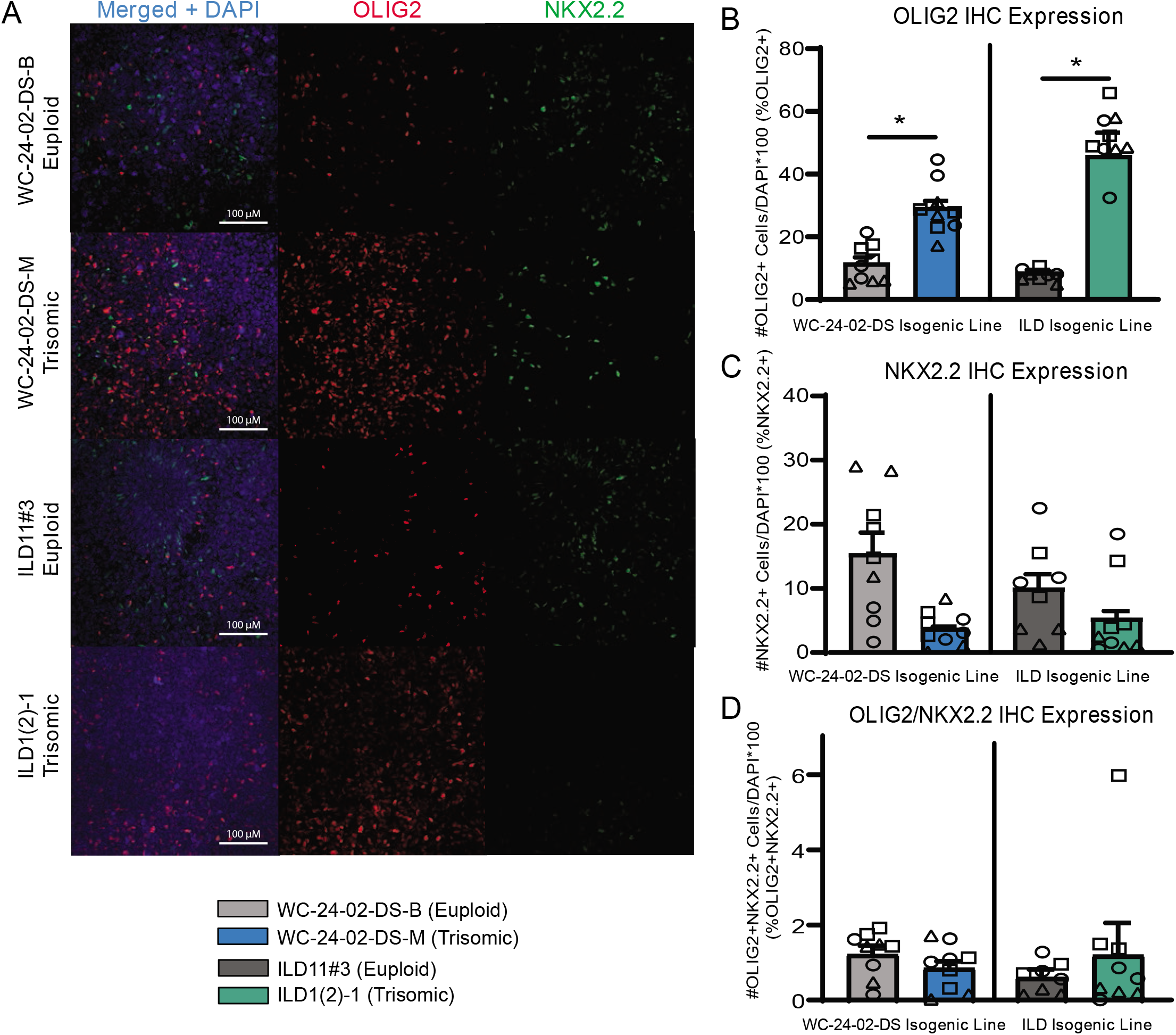
Rostral patterned tNPCs show an altered response to SHH activation. **A)** Both isogenic pairs of cells lines (WC-24-02-DS and ILD) differentiated with Wnt-C59 express OLIG2 and NKX2.2 in tNPCs after four days of treatment with SAG, a SHH pathway agonist. **B)** In both isogenic pairs, the trisomic line (WC^ts^ and ILD^ts^) has a significantly higher percentage of OLIG2+ cells in the culture. **C)** In both isogenic pairs, the trisomic line (WC^ts^ and ILD^ts^) show a trend for a lower percentage of NKX2.2+ cells in the culture. **D)** There is no difference in percentage of cells co-expressing OLIG2+ and NKX2.2+ between euploid and trisomic lines in either isogenic pair, n=3,3 independent differentiation experiments for all IHC measurements with values for individual wells overlaid. Data is shown as mean ± SEM. * p-value < 0.05 with specific values in text.

### Core DEX genes and patterning-specific differences in trisomic NPCs

While treatment with SAG resulted in expression of OLIG2 and NKX2.2 in both the RA and Wnt-C59 conditions, the magnitude of expression was not the same. In the Wnt-C59 condition we observed a general decrease in expression of NKX2.2 in all cell lines and of OLIG2 in the euploid cell lines **(Supplemental Figure 2)** though it only reached significance in the WC^ts^ NKX2.2 (p=0.021), ILD^eu^ NKX2.2 (p=0.0074), and ILD^eu^ OLIG2 (p=0.010) conditions. This may reflect the well-described crosstalk between the Wnt and SHH pathways with Wnt signaling typically promoting the expression of components of the SHH pathway (Borycki et al., 2000; Ye et al., 2009; Wang et al., 2018) that may affect trisomic and euploid cells differently.

To further elucidate gene expression differences between euploid and trisomic tNPCs, we performed bulk RNA-seq analysis on the WC lines differentiated with Wnt-C59 or RA. In the Wnt-C59 condition, we identified a total of 4,710 DEX genes between euploid and trisomic tNPCs after 4 days of SAG treatment (1μM). Of these genes, 2,199 genes were significantly upregulated in the WC^ts^ tNPCs and 2,511 genes were significantly downregulated **(Figure 4A)**. The WC^ts^ tNPCs differentiated with RA also showed large numbers of DEX genes (n=6,238). Of these, 2,803 were significantly upregulated in the trisomic tNPCs and 3,435 were down regulated **(Figure 4B)**.

**Figure 4 –.**
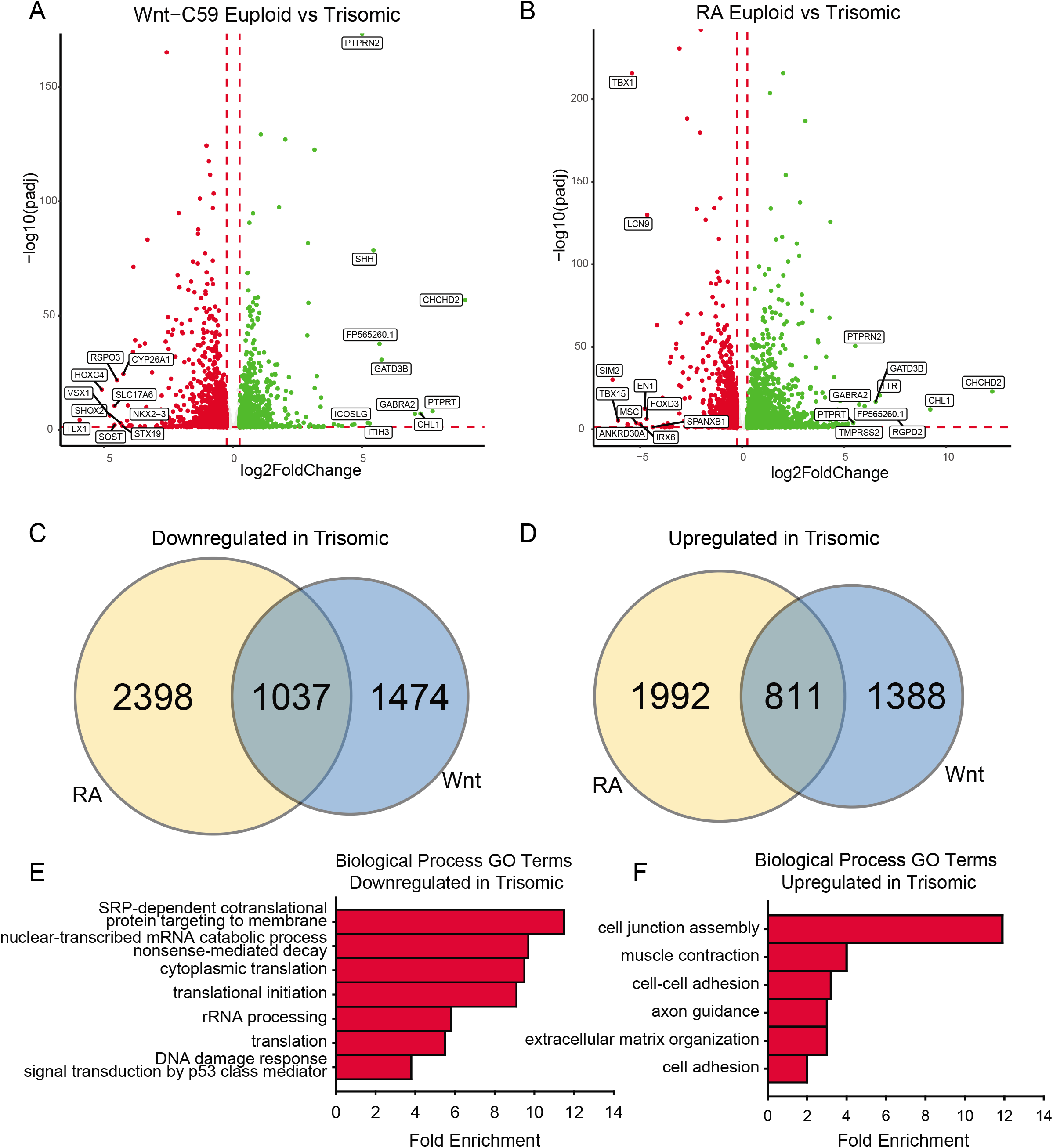
Core DEX genes and patterning-specific differences in trisomic NPCs. **A)** There are 4,709 DEX genes between euploid (WC^eu^) and trisomic (WC^ts^) tNPCs when differentiated with Wnt-C59, 2,199 upregulated in trisomic cells and 2,510 downregulated. **B)** There are 6,238 DEX genes between euploid (WC^eu^) and trisomic (WC^ts^) tNPCs when differentiated with RA, 2,803 upregulated in trisomic cells and 3,435 downregulated. **C)** Of the genes downregulated in the trisomic cultures, only 21.1% (1037) are in common between the Wnt-C59 and RA condition. **D)** Of the genes upregulated in the trisomic cultures, only 19.4% (811) are in common between the Wnt-C59 and RA condition. **E)** GO analysis identifies biological processes involving translation as significantly enriched in the genes that are downregulated in trisomic cells in both conditions. **F)** GO analysis identifies biological processes involving the extracellular matrix and cytoskeleton as significantly enriched in the genes that are upregulated in trisomic cells in both conditions.

Despite these high numbers of dysregulated genes, only a small percentage of them were consistently disturbed in both patterning conditions. In trisomic tNPCs, 1,037 (21.1%) of the downregulated DEX genes **(Figure 4C)** and 811 (19.4%) of the upregulated genes **(Figure 4D)** were common in both patterning conditions. The remaining DEX genes were only altered in one of the conditions but not the other. This large percentage of condition-specific DEX genes indicates that trisomic tNPCs are differentially affected by the spatial patterning and the different signaling pathways that are at play regionally during nervous system development. We reasoned that the dysregulated genes common in both conditions may represent a core trisomic molecular mechanism. GO analysis of the common downregulated DEX gene set identified mRNA processing and translation as an altered biological pathway **(Figure 4E)**. Specifically, *cytoplasmic translation* (GO:0002181), *translation initiation* (GO:0070992), and *rRNA processing* (GO:0006364) were significantly enriched. When considering just the common upregulated DEX gene set, GO analysis identified extracellular matrix components and cytoskeletal components as biological processes that were significantly enriched **(Figure 4F)**. These GO terms included *cell junction assembly* (GO:0034329), *cell-cell adhesion* (GO:0098609), and *extracellular matrix organization* (GO:0030198). As these processes are not dependent on regional patterning, they may reflect general processes that are perturbed in trisomic cells. Indeed, signs of repression of translational machinery were observed in monocytes and T cells collected from people with DS (Sullivan et al., 2016) supporting this hypothesis. These findings could be important for understanding general intrinsic cellular differences that may contribute to phenotypes in DS that are not region- or cell type-specific.

### SHH pathway dysregulation in trisomic tNPCs

To better understand transcriptional changes in trisomic tNPCs as they transition towards a ventral and glial fate, we used a likelihood ratio test to identify the top 1,000 DEX genes in the combination of differentiation day and genotype **(Supplemental Table 2)**. For this analysis and downstream experiments, we focused on cells cultured in the Wnt-C59 condition because of the magnitude of changes observed in OLIG2 and NKX2.2 protein expression in the brain-like tNPCs **(Figure 3)** and because Wnt-C59 patterning consistently produces a more homogenous pool of NPCs **(Figure 1F,G)**, reducing confounding transcriptomic noise from non-NPC cells in the culture. Finally, since we have shown that regional patterning strongly affects the gene expression dysregulation in the trisomic cells, we sought to assess a population of progenitors with brain-like characteristics that most closely correspond to the cellular deficits we have reported in DS white matter.

The genes identified by the likelihood ratio test model were grouped using hierarchal clustering **(Figure 5A)**. Ten clusters were chosen to represent the ten different possible patterns of gene expression between genotypes: a) same expression in NPCs (day 8) but different expression in tNPCs (day 12); b) different expression in NPCs but the same expression in tNPCs; c) different expression at both time points with increasing expression; d) different expression at both time points with decreasing expression; and e) different expression in both NPCs and tNPCs where one genotype increases and the other decreases. Finally, for each of the conditions listed, each genotype could be the one that is upregulated compared to the other leading to the total of ten patterns of gene expression. An eigenvector (PC1) based on principal component analysis (PCA) was used to identify whether differentiation day or genotype was primarily driving expression in each cluster by correlating PC1 to differentiation day **(Figure 5B, Supplemental Figure 3A,B)**. This analysis identified that the expression pattern of genes in clusters 1 (PC1=95.248%, r=-0.94445), 2 (PC1=93.557%, r=0.955467), 7 (PC1=93.708%, r=0.933868), and 8 (PC1=88.52%, r=0.883743) are primarily driven by differentiation day and may explain changes in expression observed in trisomic tNPCs. These four clusters also contain the majority (836) of the top 1,000 DEX genes.

**Figure 5 –.**
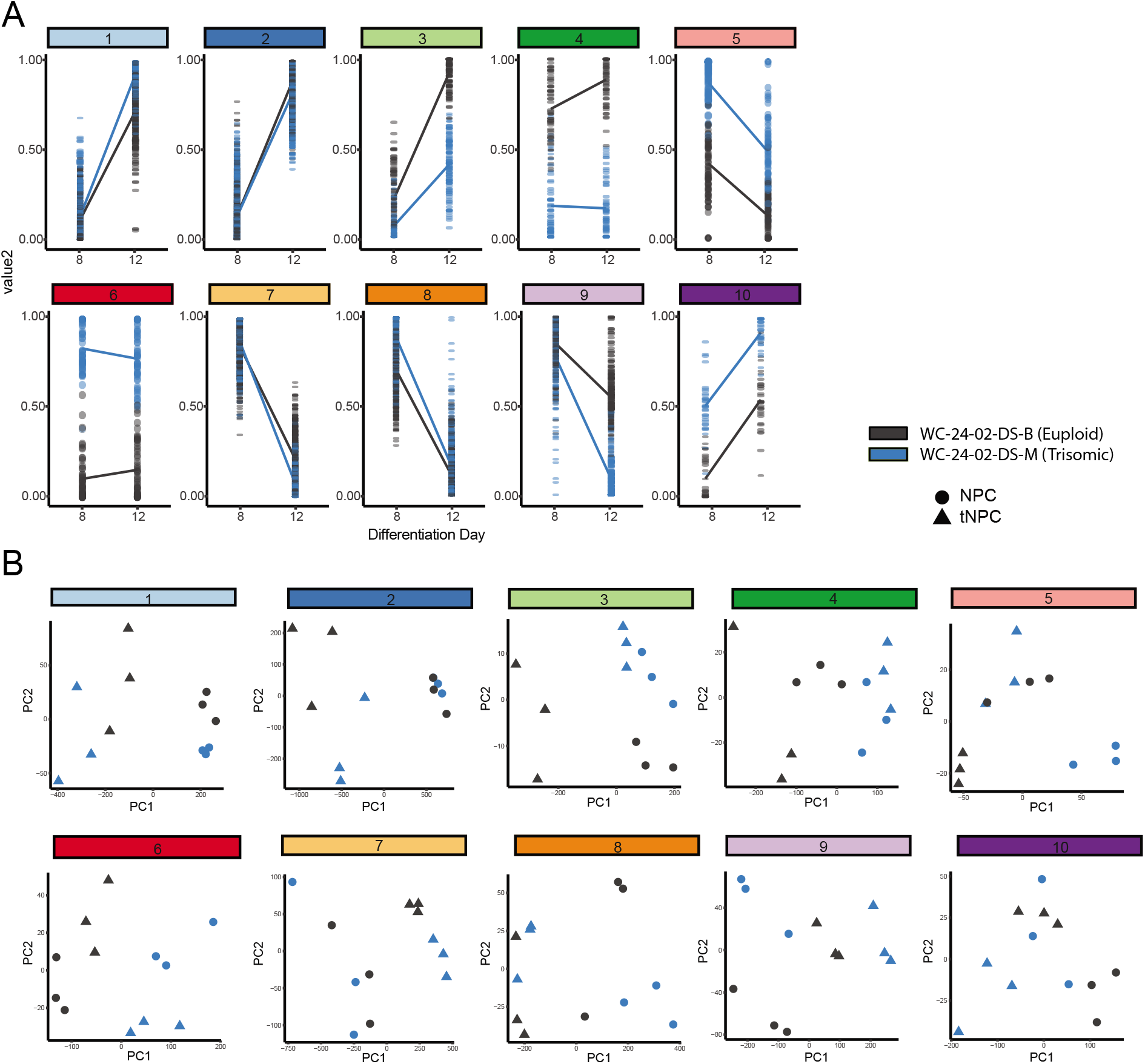
Hierarchal clustering and PCA identify gene clusters driving differentiation. **A)** Hierarchal clustering of the 1,000 DEX genes expressed during the transition of the euploid (WC^eu^) and trisomic (WC^ts^) cells from NPCs to tNPCs identifies 10 different patterns of gene expression. **B)** Principal component analysis identifies whether the primary driver of the expression pattern in each cluster is due to differentiation day or genotype by correlating PC1 with either differentiation day.

Next, the relationship between the genes was assessed using a pairwise correlation network **(Figure 6A)**. The closely correlated genes at the center of the network are predominantly from clusters 1, 2, 7, and 8, again indicating that these clusters primarily drive the differentiation of NPCs to tNPCs. Intriguingly, DEX genes that are triplicated on HSA21 are mainly located on the periphery of the pairwise correlation network **(Supplemental Figure 3C)** suggesting that they are less closely interconnected to the other 1,000 genes and perhaps represent the external drivers of the core differentiation network.

**Figure 6 -.**
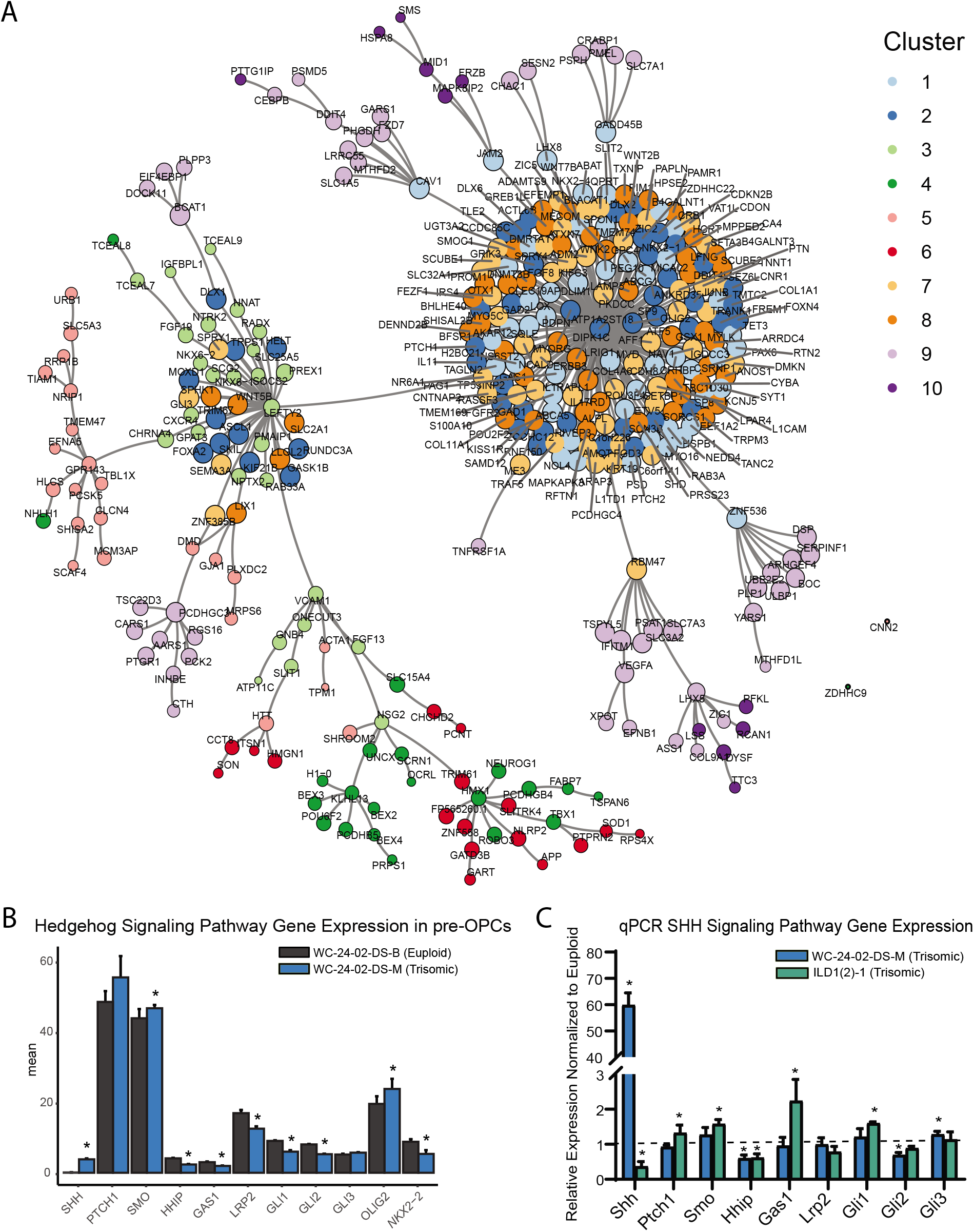
SHH pathway dysregulation in trisomic NPCs. **A)** Pairwise correlation analysis between the 1,000 DEX genes shows that gene clusters whose expression is primarily driven by differentiation day are tightly correlated in the center of the network. **B)** Closer examination of genes in the RNA-seq dataset shows that there are significant differences in SHH signaling pathway expression in the trisomic tNPCs. **C)** qRT-PCR validation of the SHH pathway genes from the RNA-seq dataset show dysregulation in both trisomic lines (WC^ts^ and ILD^ts^) of differentiated tNPCs compared to their euploid controls (WC^eu^ and ILD^eu^). n=3-5 independently cultured wells. Data is shown as mean ± SD. * p-value < 0.05 with specific values in text.

GO analysis was then performed to identify biological process and pathways that are enriched in the 10 gene clusters. For the clusters that appear to be driven mainly by differentiation, GO analysis identified a series of enriched signaling pathways **(Supplemental Figure 3D)**, including the hedgehog signaling pathway as the most enriched pathway. The differences in expression of the *OLIG2* and *NKX2.2* genes we observed in our trisomic tNPCs **(Figure 3)** are therefore consistent with perturbations in the SHH pathway. While differences in expression pattern in clusters 1, 2, 7, and 8 were primarily driven by differentiation day, close inspection of the genes in these clusters showed that genes within the hedgehog pathway were dysregulated in the trisomic cells. **(Figure 6B)**. In particular, *SHH* and *SMO* were upregulated and *GAS1, HHIP, LRP2, GLI1*, and *GLI2* were all downregulated in trisomic tNPCs. Thus, similar to previous work in the developing cerebellum of mouse models of DS (Roper et al., 2006), human trisomic NPCs patterned towards rostral brain-like fate (in the Wnt-C59 condition) may have an altered response to SHH signaling.

Since the RNA-seq analysis was performed only on the WC isogenic cells, we sought to validate the findings using qRT-PCR in both isogenic lines **(Figure 6C)**. In the WC cells, we found that *SHH* and *GLI3* were significantly upregulated in the trisomic tNPCs (p=1.2×10^−4^, 0.013) and that *HHIP* and *GLI2* were significantly downregulated (p=0.0054, 0.0065). We found that components of the SHH pathway are also dysregulated in the ILD tNPCs. *PTCH1, SMO*, and *GAS1* were all significantly upregulated in the trisomic (ILD^ts^) line compared to euploid (p=0.023, 1.3×10^−4^, 0.0022) and *SHH* and *HHIP* were significantly downregulated (p=0.021, p=0.0061).). Intriguingly, qRT-PCR of tNPCs differentiated with RA also showed dysregulation of genes in the SHH pathway in both isogenic lines. In the WC line, we found that *SHH* was significantly upregulated (p=7.1×10^−4^) and *PTCH1, HHIP, GLI1*, and *GLI2* were significantly downregulated (p=9.3×10^−5^, 0.0063, 0.0072, and 0.0050 respectively). In the ILD line, *SMO* was significantly upregulated (p=0.014) and *PTCH1* and *GLI2* were significantly downregulated (p=0.048 and p=0.010) **(Supplemental Figure 4)**. Taken together, even though the exact genes differ between isogenic lines and conditions, these results show that trisomic tNPCs derived from two different donors exhibited profound alterations in gene expression of SHH pathway genes during NPC differentiation.

### Increasing SAG concentration normalizes expression of trisomic OLIG2 and NKX2.2

The combined evidence of altered hedgehog signaling from the RNA-seq and qRT-PCR validation experiments indicated that the misexpression of OLIG2 (increased) and NKX2.2 (decreased) during trisomic NPC differentiation may be due to altered responsiveness of the SHH pathway. To test this, we modulated hedgehog signaling by increasing the concentration of SAG during the four days of tNPC differentiation from the standard 1 μM concentration (Douvaras and Fossati, 2015) to 2 μM and 4 μM. When the concentration of SAG was increased, the expression of OLIG2 and NKX2.2 in the trisomic tNPCs changed to approximate the levels found in the euploid cultures at 1μM SAG **(Figure 7A)**. For example, while 11.4% (± 2.9%) of the WC^eu^ cells expressed OLIG2 compared to 29.2% (± 3.4%) of WC^ts^ cells at 1 μM SAG (p=0.04), expression of OLIG2 in the trisomic cultures dropped to 23.0% (± 4.7%) with 2 μM SAG (p=0.23) and further to 10.2% (± 0.8%) with 4 μM SAG (p=0.99). Moreover, OLIG2 expression in the 4 μM SAG trisomic tNPCs was significantly decreased compared to 1 μM SAG (p=0.029).

**Figure 7 –.**
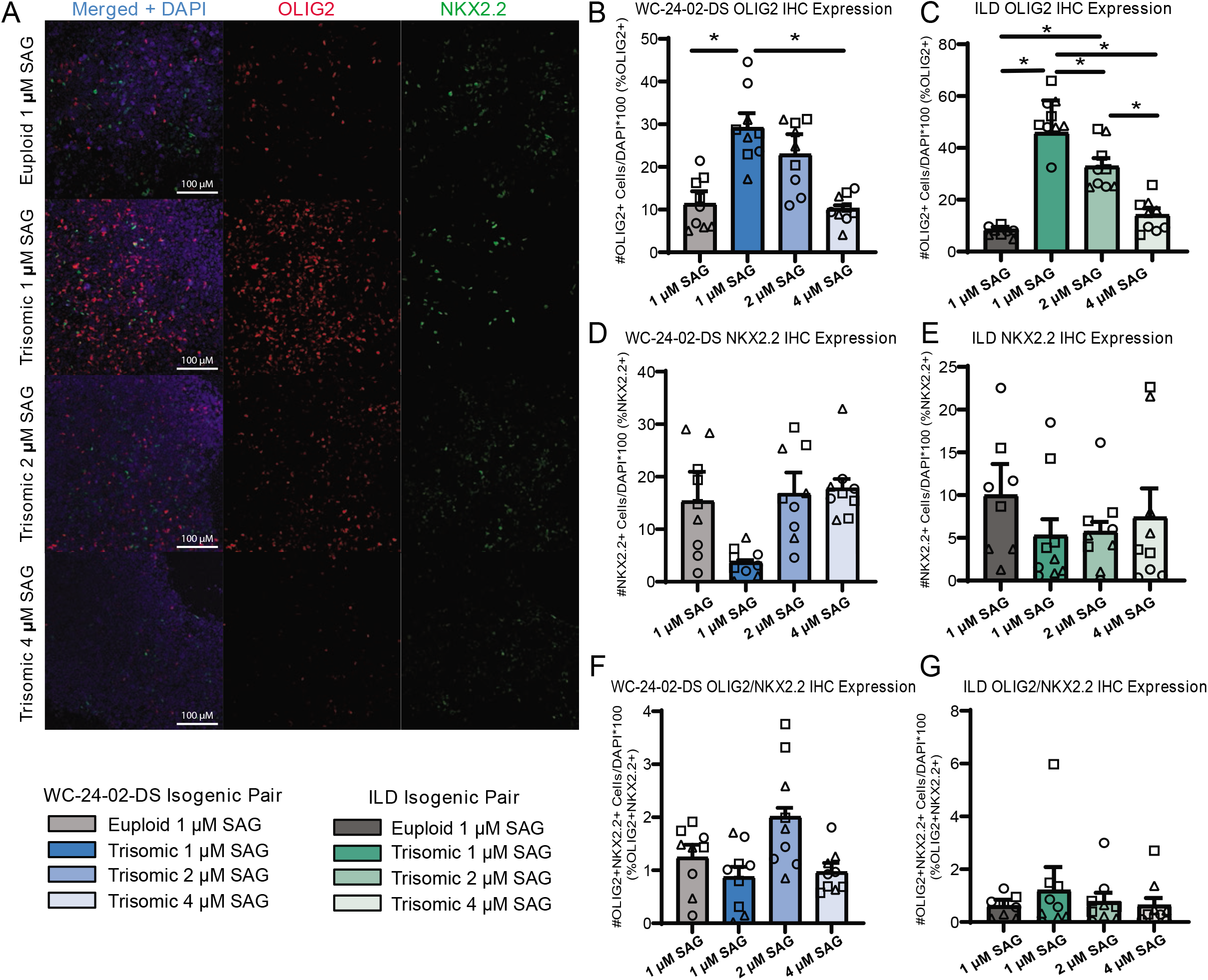
Increasing SAG concentration normalizes expression of trisomic OLIG2 and NKX2.2. **A)** Increasing the concentration of SAG in the trisomic cultures from 1 μM to 2 μM and 4 μM changes the expression level of OLIG2 and NKX2.2 from their baseline expression in trisomic cultures. **B,C)** Increasing the concentration of SAG decreases the percentage of cells expressing Olig2 in the WC^ts^ and ILD^ts^ trisomic lines. **D,E)** Increasing the concentration of SAG increases the percentage of cells expressing Nkx2.2 in the WC^ts^ and ILD^ts^ trisomic lines. **F,G)** Increasing the concentration of SAG does not change the percentage of trisomic cells co-expressing both OLIG2 and NKX2.2, n=3,3 independent differentiation experiments for all IHC measurements with values for individual wells overlaid. Data is shown as mean ± SEM. * p-value < 0.05 with specific values in text.

This pattern of expression was repeated in the SAG treatments of the ILD isogenic line **(Figure 7C)**. At 1 μM SAG, only 7.8% (± 0.7%) of the ILD^eu^ tNPCs expressed OLIG2 while 51.0% (± 2.8%) of the ILD^ts^ tNPCs expressed OLIG2 (p=2.0×10^−6^). Expression of OLIG2 in ILD^ts^ tNPCs dropped to 32.8% (± 3.2%) with 2 μM SAG (p=0.0010) and further to 14.1% (± 2.6%) with 4 μM SAG (p=0.54). Finally, there was also a significant decrease in OLIG2 expression between the 2 μM and 4 μM trisomic tNPC cultures (p=0.0068).

While OLIG2 expression decreased in the trisomic tNPCs with increasing SAG concentration, we found that this treatment led to a corresponding increase in NKX2.2 protein expression in the WC^ts^ cells **(Figure 7D).** At baseline (1 μM SAG), NKX2.2 was expressed in 15.4% (± 5.6%) of the WC^eu^ cells and only 3.7% (± 0.4%) of the WC^ts^ cells. Expression of NKX2.2 in WC^ts^ cells increased to 16.8% (± 4.0%) with 2 μM SAG and 17.8% (± 1.8%) with 4 μM SAG. While there are no significant differences between any condition, the initial trend of decreased NKX2.2 expression in the trisomic cells at 1 μM SAG disappeared when the concentration increased in trisomic cultures.

Again, this pattern of NKX2.2 expression was repeated in the ILD isogenic line **(Figure 7E)**. At 1 μM SAG, ILD^eu^ cells expressed NKX2.2 in 10.0% (± 3.7%) of the culture while only 5.3% (± 1.9%) of the ILD^ts^ cells expressed NKX2.2. Similar to the WC isogenic pair, while there are no significant differences between condition, in the presence of 2 μM SAG, the ILD^ts^ cells expressed NKX2.2 in 5.7% (± 1.2%) of the culture and this increased to 7.4% (± 3.4%) with 4 μM SAG.

The percentage of cells co-expressing OLIG2 and NKX2.2 was minimal, reflecting the early stage of NPC transition to the OL lineage. Similar to the initial 1 μM treatment, there was no significant difference in percentage of cells co-expressing OLIG2 and NKX2.2 between euploid and trisomic lines in either isogenic pair when the concentration of SAG was increased **(Figure 7F,G; Supplemental Table 1)**. This may reflect that these data represent an early differentiation time point with four days of treatment with SAG.

## Discussion

In this study, we used multiple human isogenic iPSC lines and two different spatial programming protocols to identify early-stage alterations in NPC differentiation that may underlie white matter changes in people with DS. OPCs differentiate from NPCs and commit to OL lineage fate upon activation of the SHH pathway driving the expression of the transcription factors OLIG2 and NKX2.2. We found changes in the level of expression of both these transcription factors in trisomic cells after only four days of SHH pathway activation. However, the dysregulation of these transcription factors in the trisomic cells is entirely dependent on differentiation condition and the regional patterning of the NPCs.

When the NPCs were patterned with RA toward a caudal spinal cord-like fate, we detected no change in OLIG2 or NKX2.2 expression in trisomic compared to euploid tNPCs following SAG treatment. However, when Wnt-C59 was used to pattern the NPCs towards a rostral brain-like fate, we found a decrease in NKX2.2 expression and an increase in OLIG2 expression in the trisomic tNPCs following SAG treatment. Nonetheless, the changes in gene expression between RA and Wnt-C59 are not restricted to *OLIG2* and *NKX2.2*. We identified thousands of genes that are dysregulated in trisomic cells between conditions, with only a minority of the genes dysregulated in both conditions. Similar to our findings in different human brain regions (Olmos-Serrano et al., 2016) and in the Ts65Dn mouse model (Aziz et al., 2019; Olmos-Serrano et al., 2016), these differences emphasize the importance of recognizing the spatial heterogeneity in DS and that different regions may not be dysregulated in the same manner based on their developmental history and trajectory. Thus, it is important not to generalize findings from one specific cell type into broad conclusions about dysregulated genes and pathways in DS until similar findings are reported in separate models or tissue types.

Similarly, we believe a strength of the study is the use of two separate isogenic lines derived from people with DS as this both increases rigor and helps to identify genetic differences between individuals. The replication of the differing patterns of expression of OLIG2 and NKX2.2 in RA and Wnt-C59 conditions in each line strengthens the argument that these are broad phenotypes associated with NPC differentiation in DS. However, we also identified differences between the lines, including a significantly higher percentage of Wnt-C59 patterned NPCs expressing OLIG2 in the ILD^ts^ NPCs compared to the WC^ts^ (p=0.0019) NPCs, even though both trisomic lines contained significantly more OLIG2+ cells than their euploid counterparts. Similarly, although the SHH pathway is dysregulated in both WC and ILD trisomic lines, we found variation in the transcription levels of particular genes **(Figure 6)**. The consequences of these differences between isogenic lines are thus far unknown, in part because the clinical outcomes in the two individuals are unknown.

Since several lines of evidence indicate that changes in white matter development and maintenance may play a role in the ID in DS, our goal was to understand how early OPC development may be affected in the brain. We found that NPCs differentiated with Wnt-C59 express brain-like transcriptional signatures and that trisomic Wnt-C59 patterned tNPCs express more OLIG2 and less NKX2.2 four days after SAG treatment. While the overexpression of OLIG2 in this condition can be attributed to its triplication, it is important to note that triplication of *OLIG2* does not necessarily guarantee its overexpression since we found no significant increase in OLIG2+ trisomic NPCs in the RA patterning condition. RA exposure may perhaps lead to compensatory mechanisms that inhibit the triplication-induced OLIG2 overexpression. This is not surprising considering the unique transcriptional signatures promoted by RA versus Wnt-C59 treatment **(Figure 4)**. Meanwhile, our results indicate that the decrease in NKX2.2 expression is more likely due to decreased activation and responsiveness to the SHH pathway. Specifically, in experiments where increasing amounts of SAG were applied, NKX2.2 expression was increased in the trisomic tNPCs.

This impaired responsiveness to the SHH pathway has been reported before in the Ts65Dn mouse model of DS (Currier et al., 2012; Roper et al., 2006; Trazzi et al., 2011), and we show that this deficit is present in human differentiated NPCs as well. However, previous work and our transcriptome profiling indicates that this SHH pathway dysregulation and dysfunction is not due to the triplication of SHH pathway genes on HSA21. Instead, the dysregulation of the pathway may stem from several different trans-acting sources. Previous reports have indicated that the triplication of Amyloid Precursor Protein (*APP*) may drive increased expression of *PTCH1* which therefore causes the decrease the responsiveness of SHH signaling in DS (Trazzi et al., 2011). While we do observe increased expression of *PTCH1* in the trisomic ILD^ts^ tNPCs we do not in the WC^ts^ trisomic line. In addition, at least 30 genes coding for transcription factors are located on HSA21 (Antonarakis, 2017) which, when triplicated, may indirectly regulate SHH pathway gene expression. In our study, increasing the concentration of SAG applied to the cells normalized the proportion of trisomic cells expressing OLIG2 and NKX2.2 to euploid levels **(Figure 7)**. Similarly, previous work in a mouse model of DS has shown that using SAG to modulate the SHH pathway normalized the number of granule cell precursors in the cerebellum to euploid levels (Roper et al., 2006). Altogether, several lines of evidence from multiple models confirm that modulation of SHH pathway function could be therapeutic tool for some of the neurodevelopmental deficits associated with DS, especially as future work continues to improve cell-specific targeting.

While SHH pathway dysregulation may explain the associated decrease in activation and trisomic tNPC phenotype, data from our RNA-seq analyses suggest an additional possible explanation. GO analysis of the upregulated genes in both the Wnt-C59 and RA culture conditions and analysis of the hierarchal clusters containing the majority (22/30) of the HSA21 genes point to changes in cytoskeletal components **(Supplemental Figure 3E)**. The enrichment of this category in multiple conditions indicates that cytoskeletal dysregulation is an intrinsic characteristic of trisomic cells. While genes encoding cytoskeletal components do not directly affect function of the SHH pathway, resulting changes to the cytoskeleton may affect Sonic signaling. Primary cilia are formed by different cytoskeletal proteins and SHH pathway proteins are trafficked to their correct locations via cilial transport (Wheway et al., 2018). This compartmentalization allows the signaling components to be in close proximity and mediates the strength and activity of the pathway (Gigante and Caspary, 2020; Rohatgi et al., 2007; Sasai and Briscoe, 2012). If the cytoskeleton is dysregulated in DS, primary cilial function may be impaired and thereby impact SHH signaling. In fact, recent work on fibroblasts isolated from people with DS indicates that overexpression of Pericentrin, a centrosome protein that controls microtubule organization, impacts cilia formation and function in trisomic cells (Galati et al., 2018). In addition to modulating signaling via the primary cilia, cytoskeleton remodeling is essential for OL migration, the formation of the process extensions, and the generation of myelin ensheathment (Ghelman et al. 2021). Further exploring changes in the cytoskeletal system in DS and its connection to OL differentiation and maturation will be an exciting area for future work.

Finally, the downstream effect of *OLIG2* overexpression on other types of cells is unknown. Our culture is a heterogeneous pool of NPCs, not all of which may commit to the OL lineage, reflecting *in vivo* development where different proliferative niches produce multiple different types of cells. In the same proliferative regions that give rise to OPCs, OLIG2 expressing cells also differentiate into interneurons in the brain and into motor neurons in the spinal cord (Novitch et al., 2001; Petryniak et al., 2007). The increased expression of OLIG2 in our heterogeneous culture may indicate a driver for changes in cell fate *in vivo* where the NPCs that should commit to OL fate end up committing to a different fate. This potential fate switch may result in the generation of fewer OPCs and the reduced myelin seen in people with DS. The excess OLIG2 positive cells may become interneurons in the brain, supporting reports that there may be more interneurons in the brains of people with DS (Chakrabarti et al., 2010; Xu et al., 2019). Our culture reflects a very early point in NPC differentiation along the OL lineage so this interpretation is limited to initial commitment to a pre-OPC fate and may change as cells continue to develop.

While the decrease in white matter in people with DS is well described, the etiology connecting the initial transcriptional dysregulation and the decrease in myelin is not fully understood. This study identifies changes in development very early in the process of OL specification. Altered expression of the two keystone transcription factors, OLIG2 and NKX2.2 in brain-like NPCs, may explain the changes in lineage development that results in decreased myelination. We also found that this gene dysregulation and block in NPC differentiation can be normalized by increasing the concentration of a SHH pathway agonist during differentiation. These results underscore the importance of regional and cell type differences in gene expression in DS and demonstrate that modulation of SHH signaling in trisomic cells can rescue an early perturbed step in neural lineage specification in DS.

## Acknowledgements

We would like to thank Dr. Anita Bhattacharyya of the University of Wisconsin for their kind gift of the WC-24-02-DS-B and WC-24-02-DS-M isogenic iPSCs. The work was supported by funding from the National Institutes of Health, NINDS R01-NS105138 and R21-HD098542. J.A.K was supported by NINDS F31-NS118968. In addition, this work was partially supported by grant R01-AI150305 to J.M.E., the Linda Crnic Institute for Down Syndrome, and the Global Down Syndrome Foundation. We would also like to thank Dr. Ganna Bilousova and Amy Frieman from the Department of Dermatology and the Charles C. Gates Center Regenerative Medicine at the University of Colorado Anschutz Medical Campus for their work in helping generate and characterize the iPSC pair ILD11#3/ILD1(2)1. inally, we would like to thank Dr. William Tyler for reading and providing helpful feedback on the manuscript.

## Author Contributions

J.A.K, E.Z., and T.F.H conceived the idea. J.A.K. designed and performed all the experiments excepting the bioinformatics analysis which was done by Z.L. S.R. and J.C. designed and validated the ACEq counting application. A.A. and P.S. assisted with data analysis. A.L.R. and J.M.E. produced and validated the ILD11#3/ILD1(2)-1 isogenic line. J.A.K., E.Z., and T.F.H wrote the manuscript.

## Declaration of Interests

The authors declare no competing interests.

## Materials and Methods

### Generation of iPSCs from renal epithelial cells

The iPSC lines ILD11#3/ILD1(2)-1 were generated by the Linda Crnic Institute for Down Syndrome in collaboration with the Stem Cell Biobank and Disease Modeling Core at the University of Colorado Anschutz Medical Campus. Consent was obtained from donors for the Crnic Institute’s Human Trisome Project Biobank (Colorado Multiple IRB protocol 15-2170). Renal Epithelial Cells (RECs) isolated from urine specimens and expanded as previously described (Zhou et al., 2012). The generation of iPSCs from RECs was adapted from the previously published RNA-based reprogramming method (Kogut et al., 2018). Modified mRNAs encoding six human reprogramming factors, M3O (Myo-D-Oct4), SOX2, KLF4, cMYC, NANOG, and LIN28, and miRNA mimics encoding miRNA-367/302s were produced as previously described (Kogut et al., 2018). All RNA transfections were performed using Lipofectamine® RNAiMAXTM (RNAiMAX) together with Opti-MEM® I Reduced Serum Medium (Opti-MEM) (both from Thermo Fisher Scientific) in the presence of 200 ng/mL B18R as described (Kogut et al., 2018). The reprogramming of RECs consisted of nine daily transfections. Essential 8™ medium (ThermoFisher Scientific) was used starting on Day 6 of reprogramming. The iPSC-like colonies developed on Day 15-17 of reprogramming. Mature colonies were manually picked and re-plated onto a 24-well format dish pre-coated with hESC-Qualified Matrigel (Corning) according to manufacturer’s instructions in Essential 8™ medium. At passage number 5, the iPSC’s were transitioned from Essential 8™ medium to mTeSR™1 medium (StemCell Technologies) for further expansion and analysis. Upon karyotyping several iPSC clones, an isogenic clone disomic for chromosome 21 was identified. This clone was generated as a result of a spontaneous loss of a chromosome 21 during reprogramming. Short tandem repeat (STR) analysis confirmed the origin of both D21 and T21 lines from the same REC line from the same donor.

Both generated iPSC lines were mycoplasma negative and expressed the pluripotency markers Oct3/4, Nanog and TRA-1-60 as was determined by the immunofluorescence staining **(Supplemental Figure 5A)**. The following antibodies were used for analysis: Oct3/4 Alexa Fluor 546 (Santa Cruz Biotechnology sc-5279 AF546); Nanog Alexa Fluor 647 (Santa Cruz Biotechnology, sc-293121 AF647) and TRA-1-60 (Santa Cruz biotechnology sc-21705).

iPSC lines were also subjected to the short tandem repeat (STR) analysis to confirm their origin from corresponding REC lines. Cytogenetic analysis of the generated lines was performed by WiCell® using standard GTL banding (G-banding) of metaphase chromosomes. Twenty metaphase chromosome spreads were analyzed for each established line, with chromosome classification following ISCN (2016) guidelines **(Supplemental Figure 5B)**

#### hiPSC Cell Culture and Differentiation

Two isogenic iPSC lines were used in these experiments, each consisting of a trisomic line and its euploid control that are genetically identical except for the presence of a third HSA21. One pair consists of the trisomic line WC-24-02-DS-M (WC^ts^) and its euploid control WC-24-02-DS-B (WC^eu^). This line was derived from a 25-year-old woman with DS and has been thoroughly validated by the Bhattacharyya lab at the University of Wisconsin Madison and is banked at Wicell®. The other isogenic pair consists of the trisomic line ILD1(2)-1 (ILD^ts^) and its euploid control ILD11#3 (ILD^eu^) who generation and validation was described above. This line was derived from a 46-year-old woman with DS and was produced by the Espinosa Lab at the University of Colorado Boulder. Both isogenic lines were maintained in MTeSR™ Plus media (StemCell Technologies®) supplemented with penicillin/streptomycin on Matrigel® (Corning®) coated plates. They were regularly tested for mycoplasma contamination with a PCR Mycoplasma Test Kit I/C (PromoCell®).

Each iPSC line was differentiated three independent times to NPCs and tNPCs in accordance with an established OL differentiation protocol (Douvaras and Fossati, 2015) with minor changes as noted to give each isogenic line a n=3 euploid and n=3 trisomic for every analysis. Within each independent differentiation replicate, three to five independent wells of cells were cultured per genotype to produce an average value for that replicate. In some experiments, the Wnt pathway inhibitor Wnt-C59 (Tocris BioScience) was used in place of retinoic acid. In some experiments 2 μM SAG or 4 μM SAG (MilliporeSigma) was used in place of 1 μM. iPSCs used for each differentiation were between passage 18 and 45 depending on the line.

#### Immunocytochemistry

At either day 8 or day 12 of the differentiation protocol, cells were washed with 1x PBS and fixed for 20 minutes in 4% paraformaldehyde (PFA) in 1x PBS. Fixed cells were stored in 1x PBS at 4°C for up to one week prior to staining. Fixed cells were washed with 0.1% Triton® X-100 in 1x PBS (0.1% PBST) three times for 10 minutes each. Cells were then incubated in a blocking solution of 5% donkey serum in 0.1% PBST for one hour at room temperature. This was followed by incubation of the primary antibodies in the blocking buffer at 4°C overnight. After incubation, the cells were again washed three times in 0.1% PBST for 10 minutes each and incubated with secondary antibody in blocking buffer for one hour at room temperature. Following incubation, the cells were washed twice more in 0.1% PBST and once in 1x PBS for 10 minutes each before being mounted on slides with ProLong™ Gold Antifade Mountant with DAPI (ThermoFisher Scientific).

The following primary antibodies were used: rabbit anti-Pax6 (1:250, BioLegend®, 901302), rabbit anti-Olig2 (1:500, Millipore Sigma, AB9610), and mouse anti-Nkx2.2 (1:50, Developmental Studies Hybridoma Bank, 74.5A5-s). The following secondary antibodies were used: (AlexaFluor, 1:500 dilution, ThermoFisher Scientific) donkey anti-rabbit 546 (A10040) and donkey anti-mouse 488 (A21202).

#### Confocal Imaging and Cell Counting

For each experiment, three images per cultured well were captured using a Zeiss LSM 710 confocal microscope system (Carl Zeiss, GER). Z-stacks (1024×1024 resolution) of each randomly chosen region were acquired using a 20x objective lens to cover the entire depth of cells in the region. After imaging, labelled cells were then either automatically counted using a custom developed MATLAB application (ACEq) or manually counted using ImageJ Browser software. Cell counts were processed first by image, then the three images averaged to a value for the well, then the three wells were averaged for a value for each independent differentiation, and the values from each independent differentiation were averaged finally by genotype to result in a final mean ± standard error of the mean.

#### ACEq Development and Validation

ACEq’s main purpose is to provide the user with cell counts of complex 2D and 3D cultures with a large degree of customizability. The app consists of 4 tabs that allow the user to go through 4 stages of the counting process: thresholding, segmentation, overlap correction, and results. The thresholding tab consists of a binarization threshold as well as a small particle filter that will process the raw image into a binarized and size-filtered form. The segmentation tab contains the watershed value parameter that segments regions with increasing size as it is increased. The overlap tab is the site of overlap correction to detect and remove repeated cells across the z-stack. The results tab simply provides options to display and write the results to a file.

The overlap correction method consists of using a moving window to detect and remove cells present in a given slice ‘Slice A’ that are also present in the previous slice ‘Slice B’. This is done by binarizing both slices of interest after thresholding and directly comparing them to generate a logical overlap matrix. The logical overlap matrix contains true values where the two slices of interest share the same binarized pixel values. This logical overlap matrix is then array multiplied by the labeled matrix of the Slice A. Slice A’s labeled matrix is a representation of the image where values of the elements represent the cell identity. For instance, all values ‘1’ represent the regions where the first cell would be found within the image. Using the ‘mask’ is necessary as the current and previous slices cannot directly be compared since MATLAB’s segmentation and labeling algorithm is highly susceptible to noise and its serial process of labeling regions is inconsistent across slices. The logical overlap matrix allows us to bypass this limitation by converting the cellular regions on Slice B into a labeled image that can match the labels on Slice A. Area difference and centroid difference parameters were set to tune the sensitivity of this overlap correction method. If a matching cell exceeds the thresholds for the area and centroid location differences, it would not be counted as a repeating cell.

The ACEq application was first successfully utilized for images collapsed across the z-stack. This ‘2-dimensional version’ of the algorithm was utilized by a lab member experienced with hand-counting to generate values with a mean percent error of 4.75% (n = 11) **(Supplemental Table 3)**. Subsequently, a ‘3-dimensional version’ of the app was developed to quantify markers along the z-stack and correct for overlap. This version was essential to automate analysis of images with dense cellular regions. This application was also designed to be versatile as cell counting is highly dependent on user preferences. The percent error was satisfactory in this algorithm as well with a mean percent error of 4.25% (n = 9) **(Supplemental Table 4)**. The variability in percent error is attributable to the variance in image quality and cellular regions of the OLS as well as the subjective nature of the hand counted values.

#### Statistical Analysis

Either Microsoft Excel or GraphPad Prism was used for statistical analysis. An F-test was performed to compare differences in variance in PAX6+ expression at day 8 (Figure 1) and subsequently a Kolmogorov-Smirnov test was used to directly compare genotypes and conditions. A two-tailed unpaired student’s T-test was used to directly compare the isogenic euploid and trisomic pairs in the same culture conditions such as the number of tNPCs produced at Day 12 and changes in gene expression in qRT-PCR between euploid and trisomic lines (Figures 1, 2, 3, 6, S2). A two-way ANOVA with a post-hoc Tukey’s test was used to compare changes in number of tNPCs produced in different SAG concentrations (Figure 7). Results are presented in text as a mean percentage of positive cells ± the standard error of the mean. A p-value of < 0.05 was used as the cut off for significance with specific values presented in the text.

#### qRT-PCR

For gene expression studies, cDNA was produced from mRNA from cells lysed directly in the culture plate using the TaqMan® Gene Expression Cells-to-CT™ Kit (ThermoFisher Scientific). qRT-PCR was then carried out using the corresponding TaqMan™ Gene Expression Assay Kit that include exon-spanning probes for genes of interest (ThermoFisher Scientific) (product numbers listed in **Supplemental Table 5**) on a Bio-Rad CFX96 thermocycler. Analysis was performed in BioRad CFX Maestro. All values were normalized to the housekeeping genes *GAPDH* and *UBC*, and analyzed using 2–ΔΔCt method. Three to five independent wells from each genotype were used per condition and data are shown as mean ± standard deviation. A two-tailed unpaired student’s T-test was used to directly compare the isogenic euploid and trisomic pairs in the same culture conditions. A p value of < 0.05 was used as the cut off for significance with specific values presented in the text.

#### RNA sequencing and read alignment

Total RNA was isolated from three replicates for each condition of differentiated cells using RNeasy Plus Kit (Qiagen). Purified RNA was sent to Genewiz for library preparations and Illumina® Next-Generation Sequencing at 25 million reads per sample with a paired-end 150 protocol. Quality of the reads were assessed with fastqc. Paired end reads were aligned using STAR (v2.7.3a). The genome reference for STAR was generated with GRCh38 Genome Reference Consortium Human Reference 38 (hg38) downloaded from UCSC database and gene annotation downloaded from GENCODE (v38). A 2-pass alignment workflow was followed according to the manual. Genes were annotated and quantified using featureCounts (v2.0.1). Raw read counts (*c*) for a gene (*i*) of length *l* among all genes (*I*) in a sample (*j*) were converted to fragments per kilo-million (FPKM) as follows:

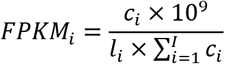

Sequencing data has been banked at NCBI Sequence Read Archive under the BioProject ID: PRJNA751601.

#### Bioinformatics analysis

Only protein coding genes were kept for downstream analyses. A p-value adjusted for multiple comparison of < 0.05 was used as the cut off for statistical significance. Principal component analysis was conducted by R function *prcomp* with FPKMs. Gene ontology (GO) analyses were performed using The Database for Annotation, Visualization and Integrated Discovery (DAVID) v6.8 (Huang et al., 2009).

Differential gene expression analysis (DEX) was conducted using *DESeq2* (Love et al., 2014). Briefly, raw read counts were used to create a DESeq dataset using the *DESeqDataSetFromMatrix* function. A manually created factor representing all combination of genotypes, differentiation conditions and differentiation days was used as the independent variable in the DESeq2 design. Then, specific pairs of comparisons were calculated using *contrast* argument in function *result*. To identify genes that show interaction between differentiation day and genotype in Wnt-C59 condition, we built another DESeq2 dataset with genotype, differentiation day and interaction of the two factors as the terms of the design formula. Then, we applied a likelihood ratio test (LRT) and genes were ranked by the LRT stat. The top 1000 genes with the highest LRT stat were kept and pairwise correlations were calculated. The correlations were converted to a distance matrix which was used for hierarchical clustering with the number of clusters set to 10, each consisting of one possible differential gene expression pattern. PCA was then conducted on the genes in each cluster and PC1 was regarded as the eigengene for each cluster.

The *igraph* R package was used to construct and represent the network of the top 1000 genes. Briefly, the pairwise correlation matrix was converted to an adjacency matrix, which is subsequently converted to a graph object by the function *graph_from_adjacency_matrix*. For genes within each cluster, only the ones degree equal to or greater than 50 were kept and unconnected vertices were removed with *delete_vertices* function. Finally, a minimum spanning tree layout was constructed with function *mst* and the graph is presented by with R package *ggnetwork*.

All plots were generated using ggplot2, unless noted otherwise.

**Supplemental Figure 1 –.**
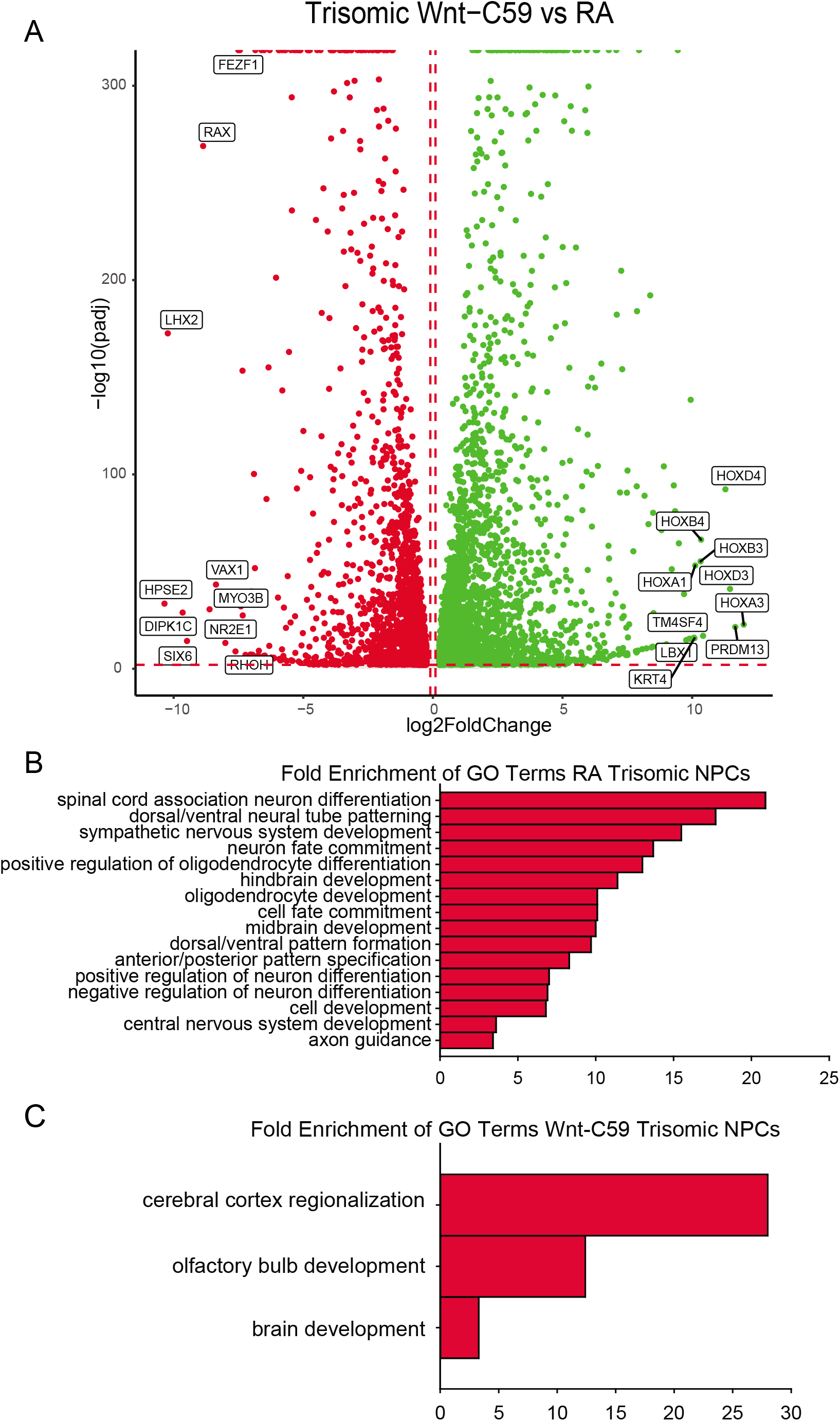
**A)** 10,115 genes are differentially expressed in trisomic NPCs differentiated with either RA or Wnt-C59. 5,169 are upregulated in the Wnt-C59 condition and 4,946 in the RA condition. **B)** GO analysis identifies biological processes significantly enriched in Wnt-C59 trisomic NPCs relating to the rostral CNS. **C)** GO analysis identifies biological processes relating the caudal CNS significantly enriched in trisomic NPCs treated with RA.

**Supplemental Figure 2 –.**
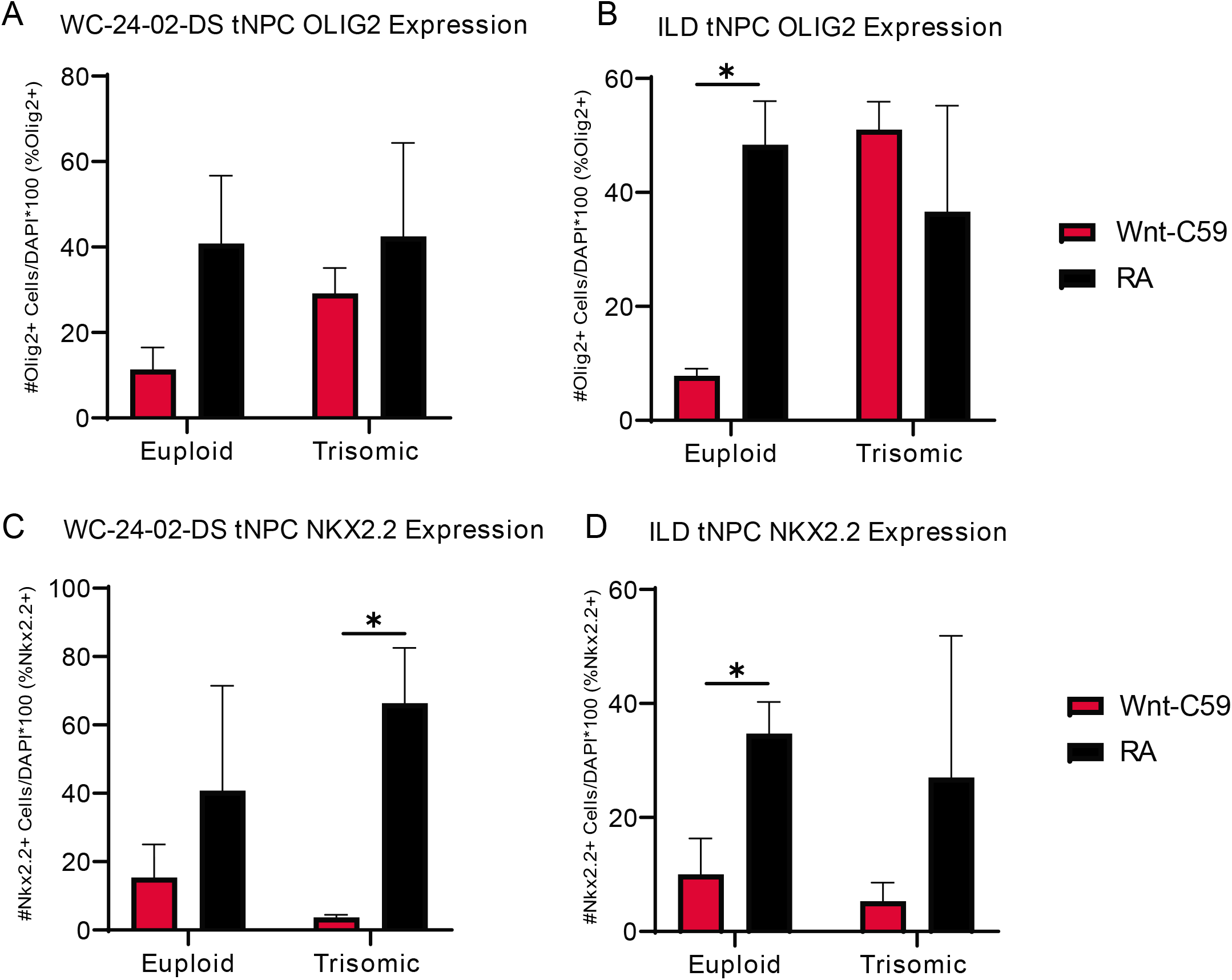
**A)** There is no difference between the RA and Wnt-C59 conditions in percentage of cells expressing OLIG2 in either genotype in the WC-24-02-DS line. **B)** In the ILD isogenic line, the Wnt-C59 euploid condition has significantly fewer OLIG2+ cells than the RA condition. **C)** In the WC-24-02-DS line, the Wnt-C59 condition generally leads to lower NKX2.2 expression than the RA condition, though it is only significant in the trisomic genotype. **D)** In the ILD line, the Wnt-C59 condition also shows less NKX2.2 expression in both genotypes though neither are significant. n=3 independent differentiation experiments, * p-value < 0.05 with specific values in text.

**Supplemental Figure 3 –.**
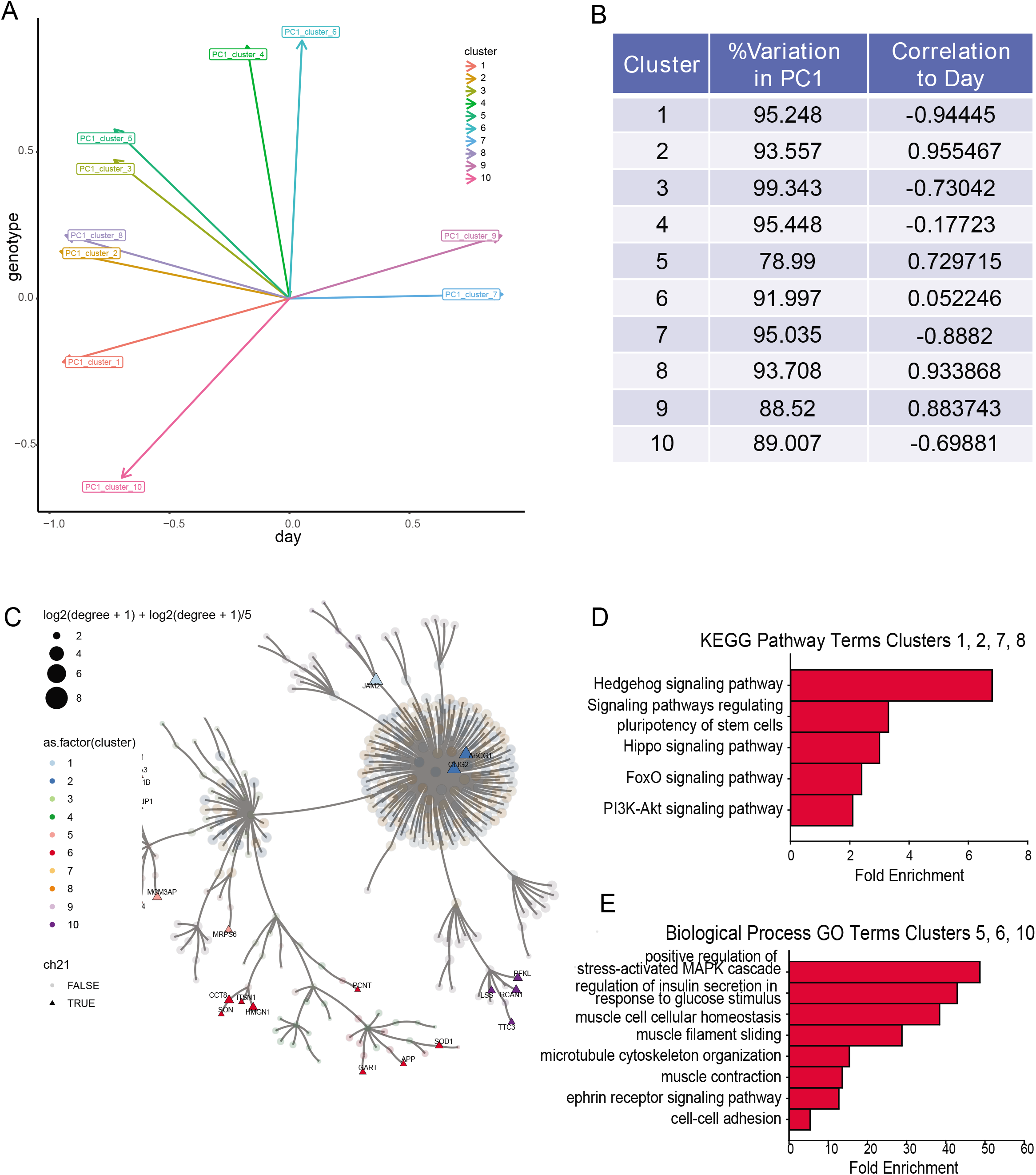
**A)** Arrows showing the correlation of principal component 1 (PC1) of each gene cluster to either day or genotype. **B)** Quantification of the percent variance explained by PC1 and its correlation to differentiation day. **C)** Location of HSA21 genes on the pairwise correlation plot. The majority of genes are peripheral way from the central node. **D)** GO analysis of the genes in clusters 1, 2, 7, and 8 identified significantly enriched KEGG pathways including hedgehog signaling as the most enriched. **E)** Gene ontology analysis of the genes in clusters 5, 6, and 10 where the genes are consistently overexpressed in the trisomic cells show multiple terms related to the cytoskeleton and extracellular matrix.

**Supplemental Figure 4 –.**
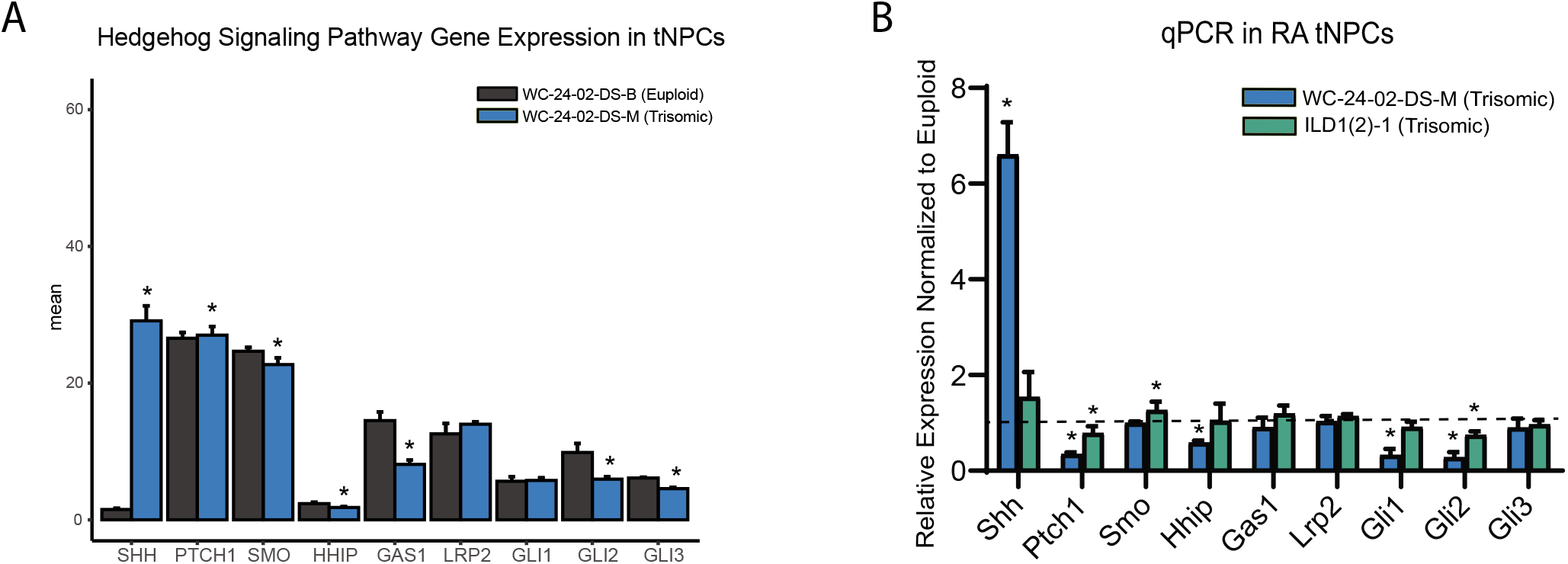
**A)** Genes in the SHH signaling pathway in our RNA-seq dataset shows that there are significant differences in expression in many of them in the trisomic tNPCs differentiated with RA. **B)** qRT-PCR validation of the SHH pathway genes from the RNA-seq dataset show dysregulation in both trisomic lines (WC^ts^ and ILD^ts^) of differentiated tNPCs compared to their euploid controls (WC^eu^ and ILD^eu^). n=3-5 independently cultured wells. * p-value < 0.05 with specific values in text.

**Supplemental Figure 5 -.**
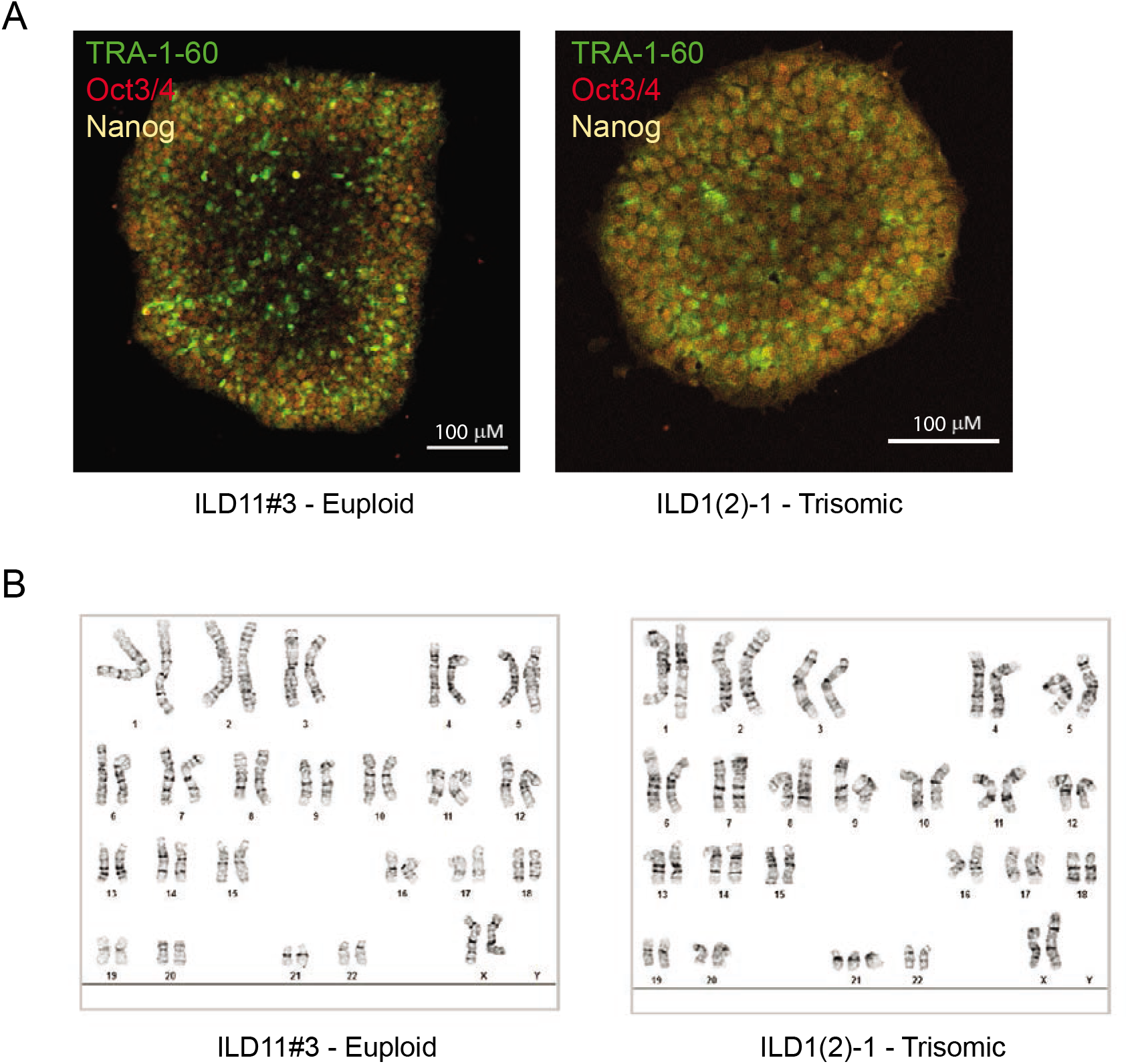
**A)** Pluripotency staining confirming expression of TRA-1-60, Oct3/4 and Nanog. **B)** Both lines were derived from the same renal epithelial cell line obtained from an individual with Down syndrome. Upon karyotyping, a euploid clone was identified (left). Short tandem repeat analysis confirmed identical origin for both lines.

**Supplemental Table 1:**
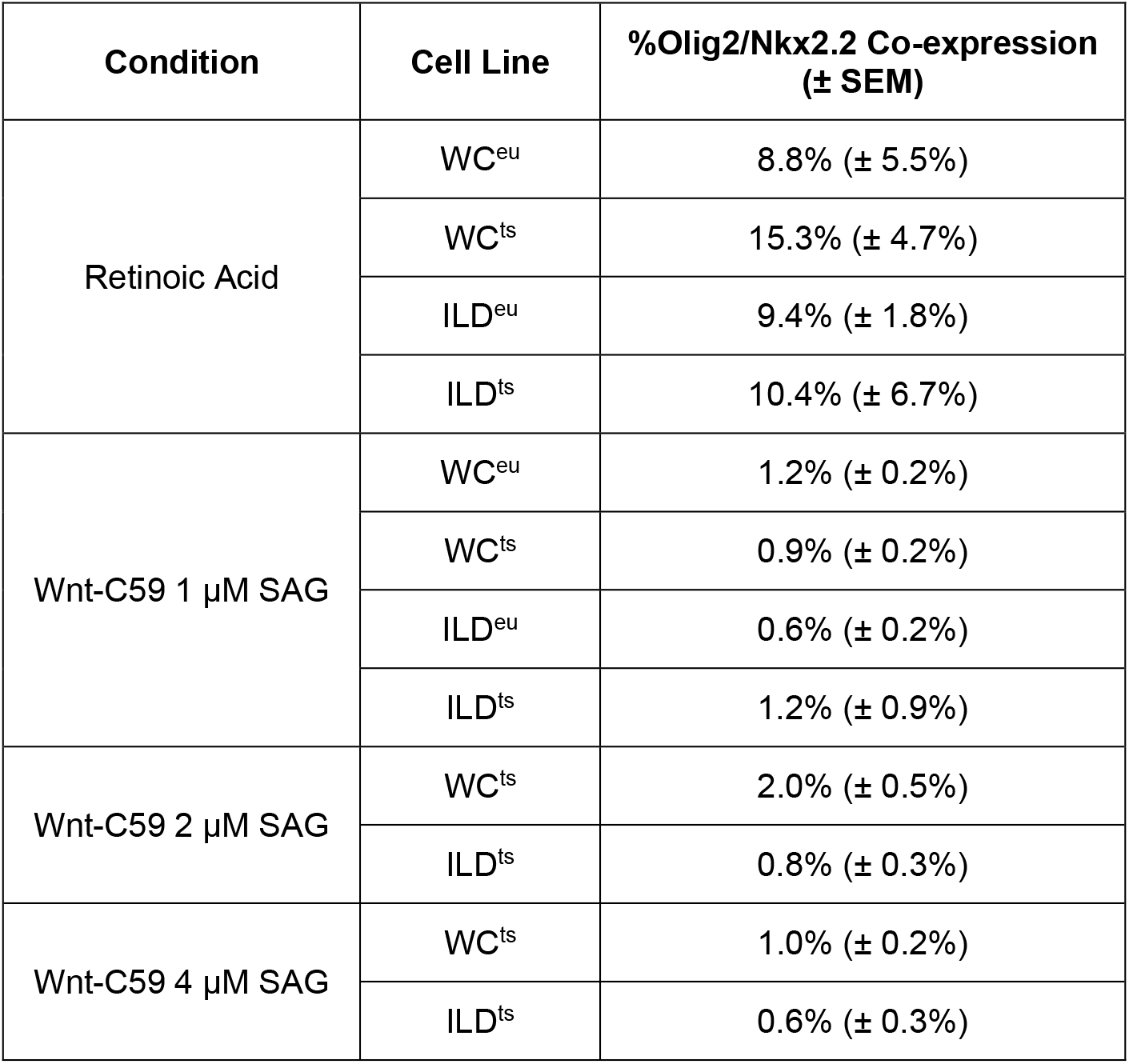
Olig2 and Nkx2.2 protein co-expression as measured by IHC in each experimental condition and iPSC line.

**Supplemental Table 2:** Top 1000 DEX genes from likelihood ratio test and assigned cluster. Please see excel file.

**Supplemental Table 3:** Computed cell counts were compared against the hand-counts for the OLIG2 and NKX2.2 stains. The DAPI stain marks too many cells and cannot be collapsed into a 2D image and will be further tested in the 3D version of the cell counting algorithm. The average error for the 2D version of the algorithm from this initial testing is 4.53%.

| Hand-Count | 2D App Cell Count | Percent Error (%) |
| --- | --- | --- |
| 121 | 121 | 0 |
| 187 | 196 | 4.8 |
| 153 | 152 | 0.65 |
| 382 | 356 | 6.8 |
| 332 | 335 | 0.9 |
| 265 | 263 | 0.7 |
| 39 | 44 | 14 |
| 381 | 402 | 5.5 |
| 177 | 168 | 5.1 |
| 316 | 297 | 6.0 |
| 335 | 317 | 5.4 |
| 303 | 317 | 4.6 |
| Average Percent Error |  | 4.53% |

**Supplemental Table 4:**
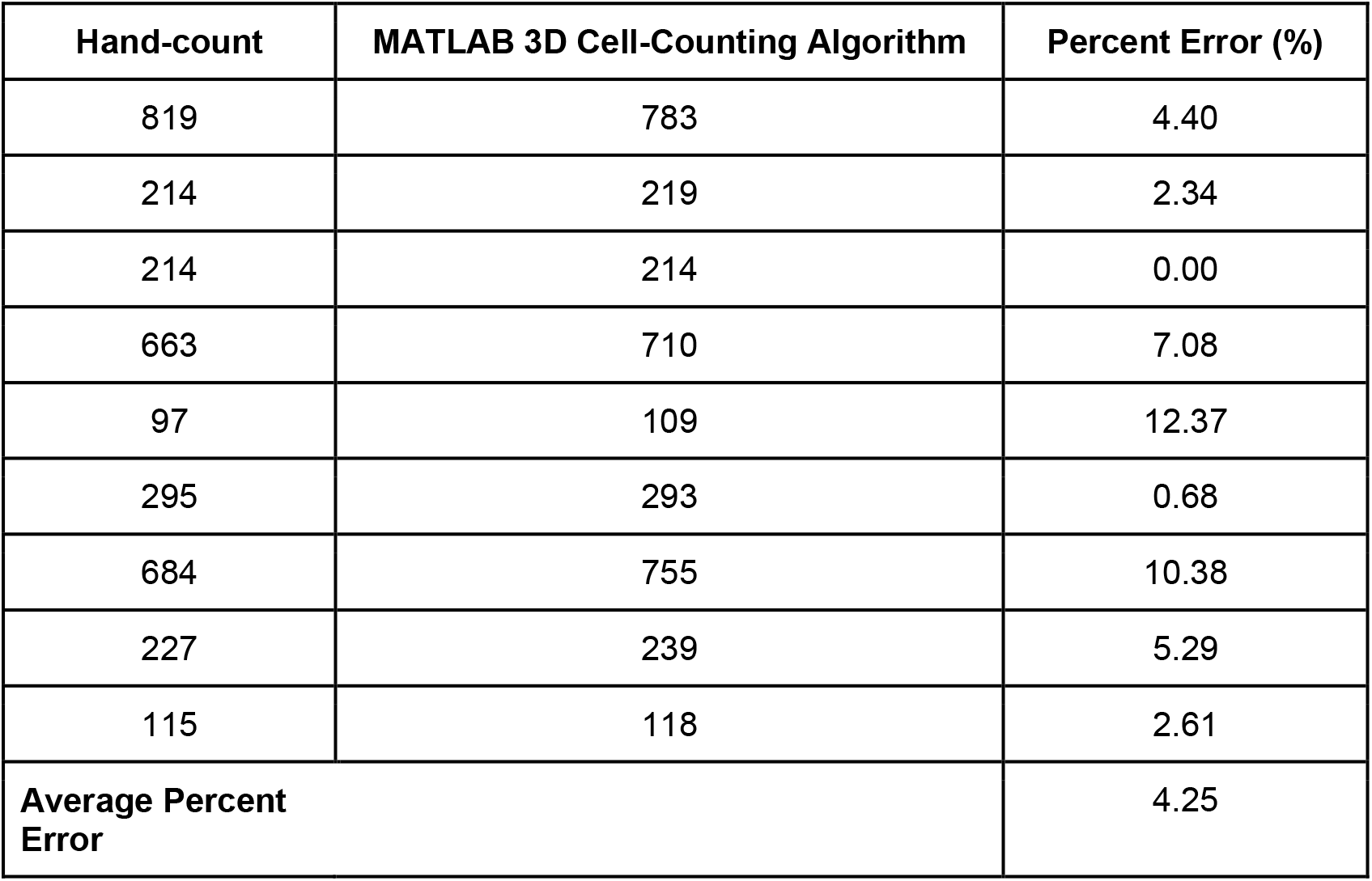
Computed cell counts were compared against the hand-counts for the DAPI, SATB2, and CTIP2 stains. The average percent error for the 3D version of the algorithm was 4.25%.

| Hand-count | MATLAB 3D Cell-Counting Algorithm | Percent Error (%) |
| --- | --- | --- |
| 819 | 783 | 4.40 |
| 214 | 219 | 2.34 |
| 214 | 214 | 0.00 |
| 663 | 710 | 7.08 |
| 97 | 109 | 12.37 |
| 295 | 293 | 0.68 |
| 684 | 755 | 10.38 |
| 227 | 239 | 5.29 |
| 115 | 118 | 2.61 |
| **Average Percent Error** |  | 4.25 |

**Supplemental Table 5.** Primers for qRT-PCR experiments. All probes are labeled with FAM dye except for *UBC* which is labeled with VIC.

| **Gene Target** | **Catalog Number** |
| --- | --- |
| *Olig2* | 4453320 |
| *Nkx2.2* | 4453320 |
| *Shh* | 4453320 |
| *Ptch1* | 4453320 |
| *Smo* | 4453320 |
| *Hhip* | 4453320 |
| *Gas1* | 4448892 |
| *Lrp2* | 4453320 |
| *Gli1* | 4453320 |
| *Gli2* | 4453320 |
| *Gli3* | 4453320 |
| *Axin2* | 4453320 |
| *GAPDH* | 4331182 |
| *UBC* | 4453320 |

## Notes

### Competing Interest Statement

The authors have declared no competing interest.

